# Reproductive isolation and structural variation among sympatric lineages of an international mosquito pest

**DOI:** 10.64898/2026.04.10.717866

**Authors:** Véronique Paris, Nancy M. Endersby-Harshman, Rahul Rane, Gunjan Pandey, Leon N. Court, Ary A. Hoffmann, Thomas L. Schmidt

## Abstract

**Background:** Reproductive isolation describes the degree to which genetic differences among populations reduce the level of neutral gene flow between them. Genomic approaches are currently showing great utility for identifying reproductive isolation between sympatric, morphologically indistinct populations, where hybridisation and introgression may yet be common. Here we use population genomic approaches and a chromosome-level reference assembly to uncover three highly differentiated, cryptic lineages of *Aedes notoscriptus*, an Australian mosquito disease vector that has invaded New Zealand (c. 1920) and California (c. 2014).

**Results:** We identified a single lineage (Noto1) that was the source of the New Zealand and Californian invasions, and this lineage was also distributed across much of Australia including the Greater Melbourne region where it was sympatric with a second lineage (Noto2). Although Victorian Noto1 and Noto2 were frequently sampled co-locally, hybrids were observed less frequently than expected under random mating, and admixture levels were low, suggestive of selection against hybrids. A bottleneck observed in Victorian Noto1 suggests that Noto1 may have recently expanded its range into the Greater Melbourne region. We also identified a massive structural variant (∼25 Mbp; ∼10% of chromosome size) in Noto2 that was segregating at intermediate frequencies across the Greater Melbourne region but was fixed in Noto1.

**Conclusion:** Together, these results identify key genetic differences between cryptic lineages of this international disease vector that warrant further investigation given potential differences in vector competence.

## 1 Background

Reproductive isolation plays a central role in the formation and maintenance of species boundaries (Ripley and Mayr 1943, Ayala and Fitch 1997). It is increasingly recognized not just as a qualitative outcome but as a measurable trait, varying along a continuum like genetic structure (Westram et al. 2022). High-resolution, genome-wide sequence data broaden our understanding of reproductive isolation by enabling researchers to detect and quantify selective pressures, reproductive barriers, and past and present gene flow beyond what morphology alone can indicate (e.g. Butlin 2010, Feder et al. 2012, Tittes and Kane 2014). Such sequence data can also identify genomic regions where specific SNPs or structural variants lead to higher or lower permeability to gene flow depending on the strength and type of selection (Turner et al. 2005, Westram et al. 2022). Reproductive isolation has a geographic component and is most clearly observed in sympatric and parapatric populations, where opportunities for hybridisation are greatest (Turelli et al. 2001). Understanding reproductive isolation has proven vital for identifying cryptic species complexes, and is important for pests like mosquitoes where species delineation is needed to manage the spread of insecticide resistance (Clarkson et al. 2014) or plan novel control interventions involving reproductive incompatibility (Hoffmann et al. 2011, Beebe et al. 2021).

In mosquitoes, reproductive isolation can operate through both pre- and post-mating mechanisms. Pre-mating isolation can arise through temporal or behavioural separation; for example, *Anopheles gambiae* and *An. coluzzii* mate at different times of day, reducing interspecific mating (Sawadogo et al. 2013). Post-mating isolation is evident in hybrid sterility and reduced fitness, as seen in F1 hybrids between *An. gambiae* and *An. arabiensis*, which often suffer reduced fertility or viability, leading to the gradual purging of introgressed genetic backgrounds (Slotman et al. 2004, Hanemaaijer et al. 2018). These mechanisms reinforce species divergence and have significant implications for both evolutionary trajectories and vector management strategies.

One mosquito species of increasing public health concern is the Australian backyard mosquito, *Aedes notoscriptus*. Originally native to mainland Australia and Tasmania, it has expanded and established populations in the Torres Strait Islands, New Zealand (Laird and Easton 1994), New Guinea, New Caledonia, Indonesia (Dobrotworsky 1965, Lee 1987, Sunahara and Mogi 2004), and more recently, California, USA (Metzger et al. 2021). In Australia, it is a competent vector of multiple arboviruses, including Ross River virus (Doggett and Russell 1997, Watson and Kay 1999) and Barmah Forest virus (Watson and Kay 1999), and is the primary vector of dog heartworm (Russell and Geary 1992). In Victoria, *Ae. notoscriptus* is specifically implicated in the transmission of *Mycobacterium ulcerans*, the causative agent of Buruli ulcer, making it one of the rare cases of a bacterium being mechanically transmitted by a mosquito species (Wallace et al. 2017, Mee et al. 2024, Paris & Buultjens et al. 2026).

Genomic analyses across Culicidae have revealed substantial differentiation among morphologically similar taxa (Pombi et al. 2014, Fontaine et al. 2015, Tennessen et al. 2021) and mitochondrial DNA studies indicate that *Ae. notoscriptus* contains at least four cryptic lineages with divergences of 2.14 – 4.36 % (Endersby et al. 2013). Correctly identifying mosquito lineages is crucial, as they may differ in ecologically and epidemiologically relevant traits (Funk et al. 2006) such as dispersal capacity, host preference (Dumas et al. 2016), vector competence (Brown et al. 2011, Karthika et al. 2018), and endosymbiont interactions (Rasgon et al. 2006, Minard et al. 2017, Guo et al. 2018). However, mitochondrial markers capture only a small fraction of the genome and discordance between mitochondrial and nuclear variation is well documented in mosquitoes (e.g. Hemmerter et al. 2009, Hlaing et al. 2009, Rašić et al. 2015, Zadra et al. 2021, Amaya Romero et al. 2024), limiting our understanding of lineage boundaries, connectivity, and adaptive divergence. Because both reproductive isolation and cryptic genomic structure can restrict gene flow within and among mosquito populations, they may have direct implications for vector control approaches such as *Wolbachia* releases and gene drives, which rely on successful mating and gene flow to sustainably spread (Hoffmann et al. 2011, Burt 2014).

Despite its growing public health importance and global invasion coverage, no geographically broad genomic studies of *Ae. notoscriptus* have yet been performed. In this study, we generate a chromosome-level reference genome assembly for *Ae. notoscriptus* and genome-wide, reduced-representation sequence data from across parts of its global range to test whether the mtDNA-defined lineages described by Endersby et al. (2013) are reflected in nuclear genomic structure, and to characterise patterns of gene flow and reproductive isolation among any nuclear lineages identified. We identify three nuclear lineages: Noto1, Noto2, and Noto3. We show that these do not align with mtDNA lineages but nevertheless exhibit reproductive isolation in two (Noto1 and Noto2) that are sympatric across the Greater Melbourne region, Australia. Our findings introduce *Ae. notoscriptus* as a compelling model for understanding reproductive isolation in sympatric lineages and have important implications for the management of this internationally invasive disease vector.

## 2 Results

### 2.1 Reference genome assembly

High-fidelity sequencing generated 52.1 Gb of PacBio HiFi data, which, together with Hi-C data, enabled the assembly of a chromosome-scale genome for *Ae. notoscriptus*. The final assembly (GenBank accession GCA_040801935.1) spans 900.4 Mb with a contig and scaffold N50 of 1.8 and 282 Mb, respectively, and comprises three primary chromosome-scale scaffolds (Chr1: 290Mb; Chr2: 282 Mb; Chr3: 205 Mb). This genome assembly also includes 305 unplaced scaffolds. Genome completeness assessed using BUSCO identified C: 95.0% [S: 89.1%, D: 5.9%], F: 0.9%, M: 4.1% (n = 3,285) for the Diptera odb10 dataset and C: 97.2% [S: 90.9%, D: 6.3%], F: 0.4%, M: 2.4% (n = 1,367) for the Insecta obb10 dataset, indicating high completeness of conserved gene content.

To test which of the three *Ae. notoscriptus* lineages the reference genome was sourced from (see 2.3), we calculated the proportion of reference and alternate alleles for mosquitoes of each lineage (Table S1). This provided clear evidence that the reference assembly was sourced from the Noto1 lineage (96.7% similarity vs <84.9–87.1% similarity).

### 2.2 Population genomic dataset

We generated genome-wide sequence data for Australian samples from Victoria (VIC), New South Wales (NSW), Queensland (QLD), the Northern Territory (NT), American samples from California (USA), and New Zealand samples from the North Island (NZ) (Figure 1, n=189 total). USA and NZ samples were from invasive populations, while the Australian samples were from the presumed native range. Our Australian sampling was concentrated in south-eastern Australia and, together with limited sampling of invasive populations, should not be interpreted as capturing the full geographic diversity of the species. Instead, our sampling was primarily focused around the Greater Melbourne region (n=155).

**Figure 1.**
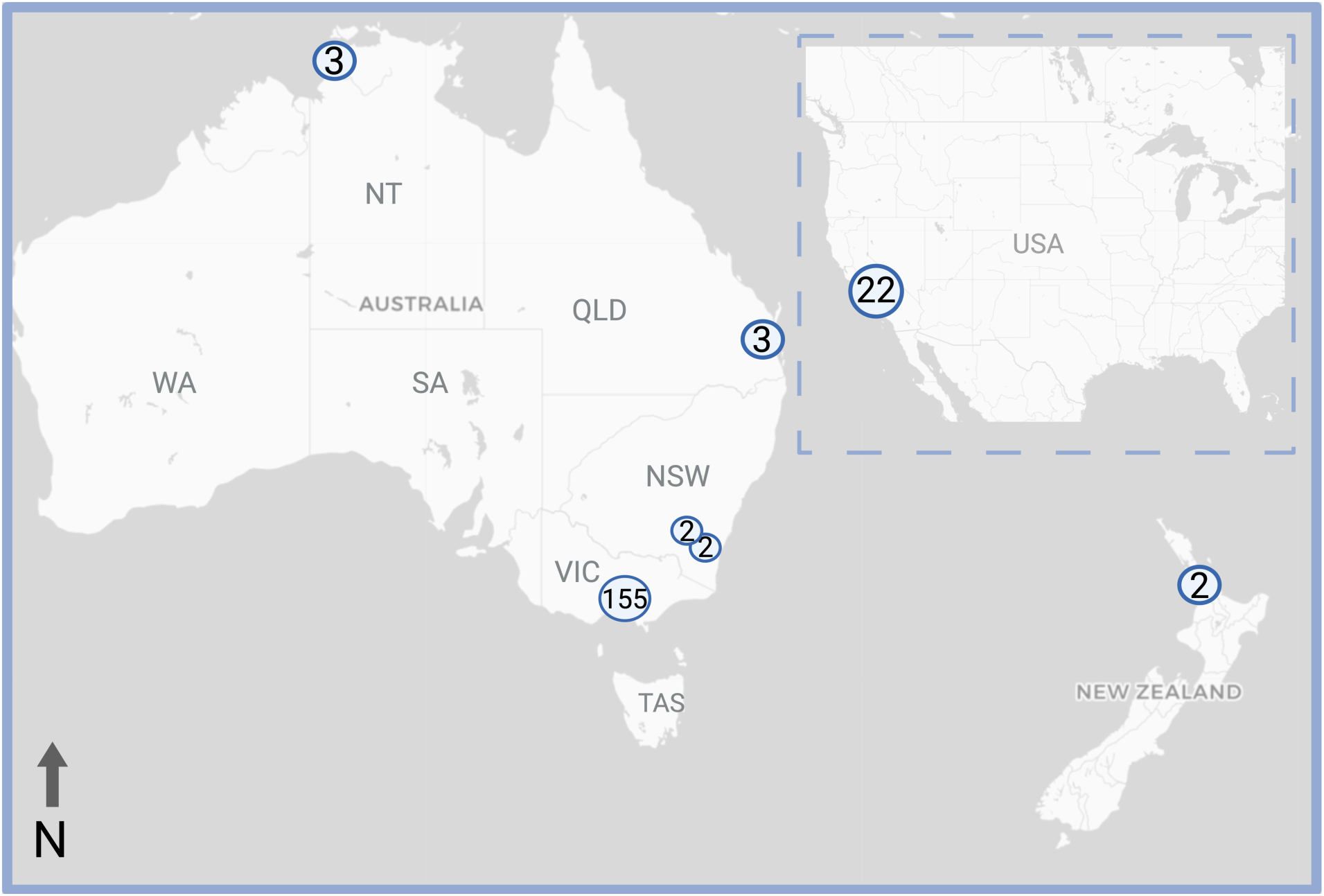
Geographic distribution of samples included in the genomic analyses in this study. Numbers in circles indicate the number of samples included in genomic analyses. Most samples were collected from south-eastern Australia, with additional samples from other parts of Australia, New Zealand and California USA (inset).

#### 2.3.1 Three deeply differentiated nuDNA lineages of Aedes notoscriptus

To explore patterns of genetic structure among individuals, we first performed a principal component analysis (PCA) on the entire global dataset (Figure S1). PCA revealed three main clusters along the first two principal components. The first cluster contained samples from the NT, which were distinct from all other populations. The second cluster contained individuals from VIC, NSW, QLD, and the invasive samples from NZ and the USA. Surprisingly, the third cluster contained another set of individuals from VIC, which were clearly distinct from the second cluster. Henceforth, we refer to the large cluster as Noto1, which contained most Victorian samples and all samples from NSW, QLD, NZ, and USA. We refer to the smaller Victorian cluster as Noto2. We refer to the NT lineage as Noto3.

To characterise clustering within Victoria, we used discriminant analysis of principal components (DAPC) implemented in R package adegenet (Jombart et al. 2010) (Figure 2A). To minimise potential bias from strong drift in invasive populations, analyses were restricted to Australian samples. Genetic clusters were first inferred using BIC-guided K-means clustering (find.clusters), and DAPC was then used to summarise and visualise differentiation among these inferred groups. The analysis supported three genetic clusters corresponding to Noto1 (n = 122), Noto2 (n = 33), and Noto3 (n = 3). We checked whether inference around lineages could be affected by differential missing data (e.g. such as from allelic dropout (Andrews et al. 2016)). Both Noto1 and Noto2 had little variation in missing data and depth, though lower depth was recorded for samples from NZ and NT (Noto3) likely due to sample degradation (Table S1). This suggests that genomic inferences should be robust around the main two lineages, Noto1 and Noto2.

**Figure 2.**
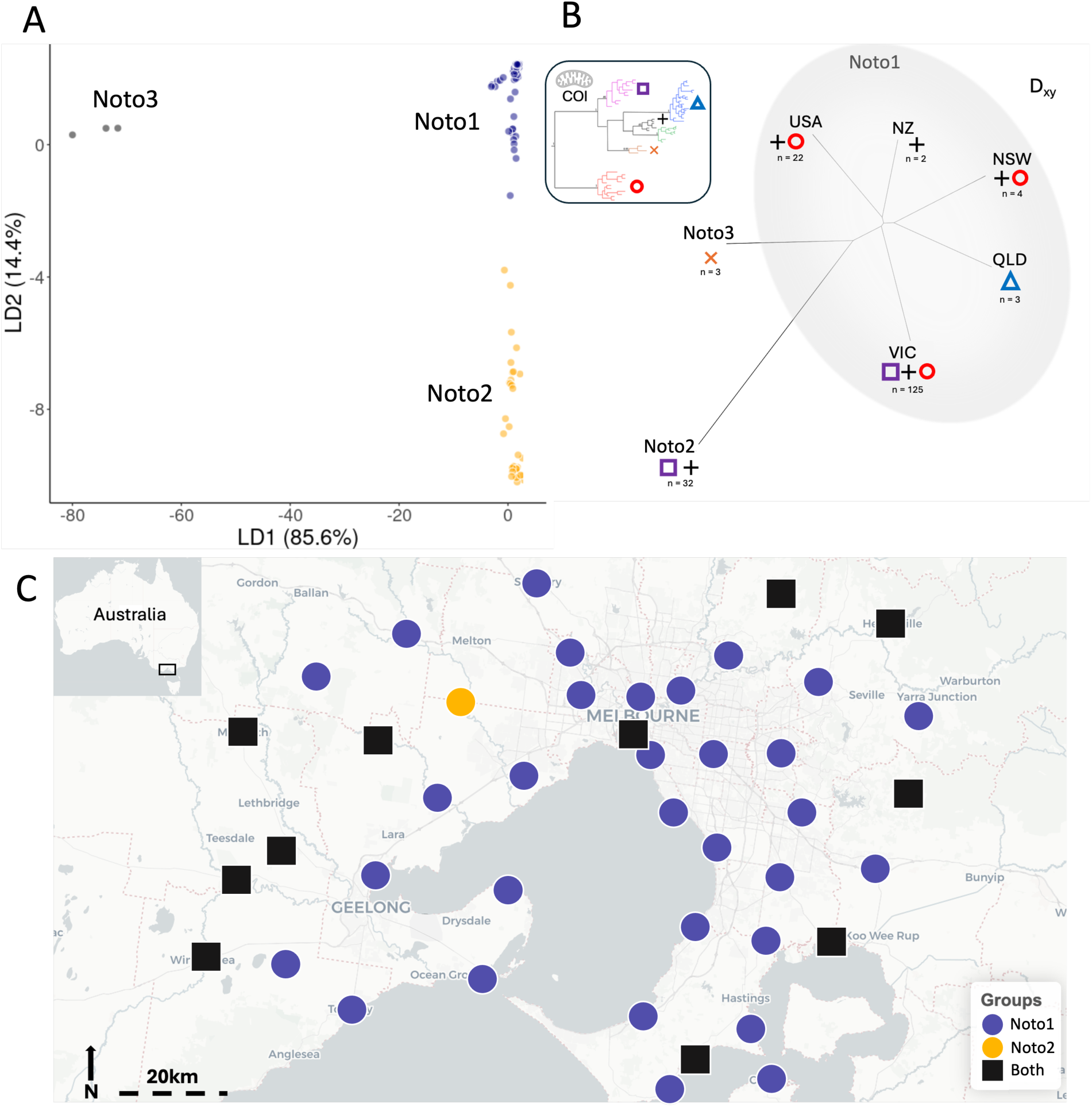
(A) Discriminant Analysis of Principal Components (DAPC) for Ae. notoscriptus across the native Australian range, using 2 PCs, following Thia (2022). Three genetic clusters were identified: Noto1(blue), Noto2 (yellow), and Noto3 (grey). Each point represents an individual, coloured according to the assigned DAPC cluster. (B) Phylogenetic network based on genome-wide d_XY_ illustrating relationships among nuclear lineages. Symbols correspond to the mitochondrial clades found in each nuclear lineage (inset), highlighting mito-nuclear discordance. Inset: COI mitochondrial haplotype tree with distinct clades indicated by colours and symbols. The full COI tree is provided in Figure S3. (C) Map of sampling locations across the Greater Melbourne region showing the geographic distribution of the two major genetic clusters. Sites where only Victorian Noto1 was sampled are shown in blue and sites where only Noto2 was sampled are shown in yellow. Locations where both clusters were sampled are indicated by black squares.

Having determined the three native range clusters, we constructed a phylogeny using genome-wide *d_XY_,* a measure of absolute divergence (Figure 2B). As *d_XY_* is minimally influenced by post-split differences in rates of genetic drift (Cruickshank and Hahn 2014), it can help differentiate between strong structure caused by long periods of reproductive isolation and structure caused by recent invasion bottlenecks. The *d_XY_* phylogeny further validated the existence of three lineages (Figure 2B), and indicated that the USA and NZ invasions were most likely sourced from the Noto1 lineage.

#### 2.3.2 Nuclear lineages are discordant with mtDNA lineages

Previous work identified five deep mtDNA lineages of *Ae. notoscriptus* (Endersby et al. 2013). We investigated whether our nuclear genomic lineages accorded with these by sequencing the mtDNA COI gene of our global dataset of individuals. Surprisingly, there was little concordance between the nuDNA and mtDNA datasets (Figure 2B, insert). While QLD and the NT each had specific mtDNA clades found only in these locations, the other mtDNA clades were distributed across the nuDNA tree, including two mtDNA clades that were found in both Noto1 and Noto2. Further sampling may uncover other mtDNA clades in QLD and the NT, which were each only represented by three samples. We analysed COI variation using an additional 237 *Ae. notoscriptus* mtDNA sequences available through GenBank (Table S2); these show that the COI clades we describe here match those of Endersby et al. 2013 (Table S2; Table S3; Figure S3). Detailed results of the mtDNA analysis can be found in the Supplementary Methods.

#### 2.3.3 Two nuclear lineages are sympatric across the Greater Melbourne region

Having established that Noto1 and Noto2 both occur in Victoria, we investigated their local geographical distributions (Figure 2C). Victorian Noto1 and Noto2 were broadly distributed across the Greater Melbourne region, with Victorian Noto1 found in more locations than Noto2. Eleven out of 43 sites (26%) contained both Victorian Noto1 and Noto2. Note that this co-distribution is likely an underestimate, as six out of 43 sites had only one sequenced mosquito, with a maximum number of five from any single site. Nevertheless, Victorian Noto1 and Noto2 were clearly both widespread, and found in central Melbourne as well as in regional areas extending in multiple directions from the city centre.

#### 2.3.4 No Wolbachia infection detected in any Aedes notoscriptus

Endosymbiotic *Wolbachia* bacteria are widespread in some *Aedes* mosquitoes (Raquin et al. 2015, Puerta-Guardo et al. 2020) but not others (Ross et al. 2020, Ross and Hoffmann 2024). *Wolbachia* infection in *Ae. notoscriptus* has been reported in some populations from Queensland (Skelton et al. 2016). The infection status of *Ae. notoscriptus* in southern Australia remains uncertain (Duchemin et al. 2017) and could vary across lineages, potentially contributing to reproductive isolation through *Wolbachia*-mediated effects on host phenotypes, such as bidirectional cytoplasmic incompatibility (Rousset et al. 1992, Stouthamer et al. 1999). We screened all individuals for the presence of *Wolbachia* using two methods, qPCR and sequence alignment to *Wolbachia* genomes. Both approaches found that every individual in our dataset was uninfected with *Wolbachia,* including the three samples from Queensland.

### 2.4 Hybridisation is rare in sympatric lineages of Aedes notoscriptus

The strong mito-nuclear discordance across *Ae. notoscriptus* (Figure 2B) could be explained by two non-mutually exclusive hypotheses. The first is that Noto1 and Noto2 are still able to hybridise, and either reciprocal backcrossing or selection against hybrids leads to the redistribution of mtDNA lineages across ’purified’ nuDNA lines (Jordal et al. 2002). The second is that the reproductive isolation between Noto1 and Noto2 has begun too recently for lineage sorting to have fixed mtDNA lineages in each nuclear line (Hudson and Coyne 2002). As Victorian Noto1 and Noto2 co-occur across the Greater Melbourne region (Figure 2C), there appears no current geographical impediment to hybridisation. To investigate the above hypotheses, we used population genetic analyses specifically capable of detecting and characterising admixture: *ADMIXTURE* (Alexander et al. 2009), *Treemix* (Pickrell and Pritchard 2012) and *fineRADstructure* (Malinsky et al. 2018).

*ADMIXTURE* showed very clear separation of the Noto1, Noto2, and Noto3 clusters, as well as the invasive USA cluster within Noto1 (Figure 3A). However, four individuals had genotypes consisting of roughly equal parts of Victorian Noto1 and Noto2, with each having >38% of each background (Figure 3A). We investigated these further by projecting the four samples onto a principal component constructed from non-admixed Victorian Noto1 and Noto2 individuals (Figure S2). The four samples were all placed between the putatively non-admixed samples, as expected if they were the product of hybridisation. Two of the four samples were placed between the Victorian Noto1 and Noto2 clusters, so that they could not be identified as predominantly of either lineage (Figure S2). Consistent with recent hybrid ancestry, the four putative hybrids also displayed elevated genome-wide heterozygosity compared with non-admixed Victorian Noto1 and Noto2 individuals (Figure 3D). When plotted against VIC2 ancestry proportions, these individuals occupied intermediate ancestry positions with increased heterozygosity relative to parental lineage individuals, consistent with expectations for recent-generation hybrids (Figure 3D).

**Figure 3.**
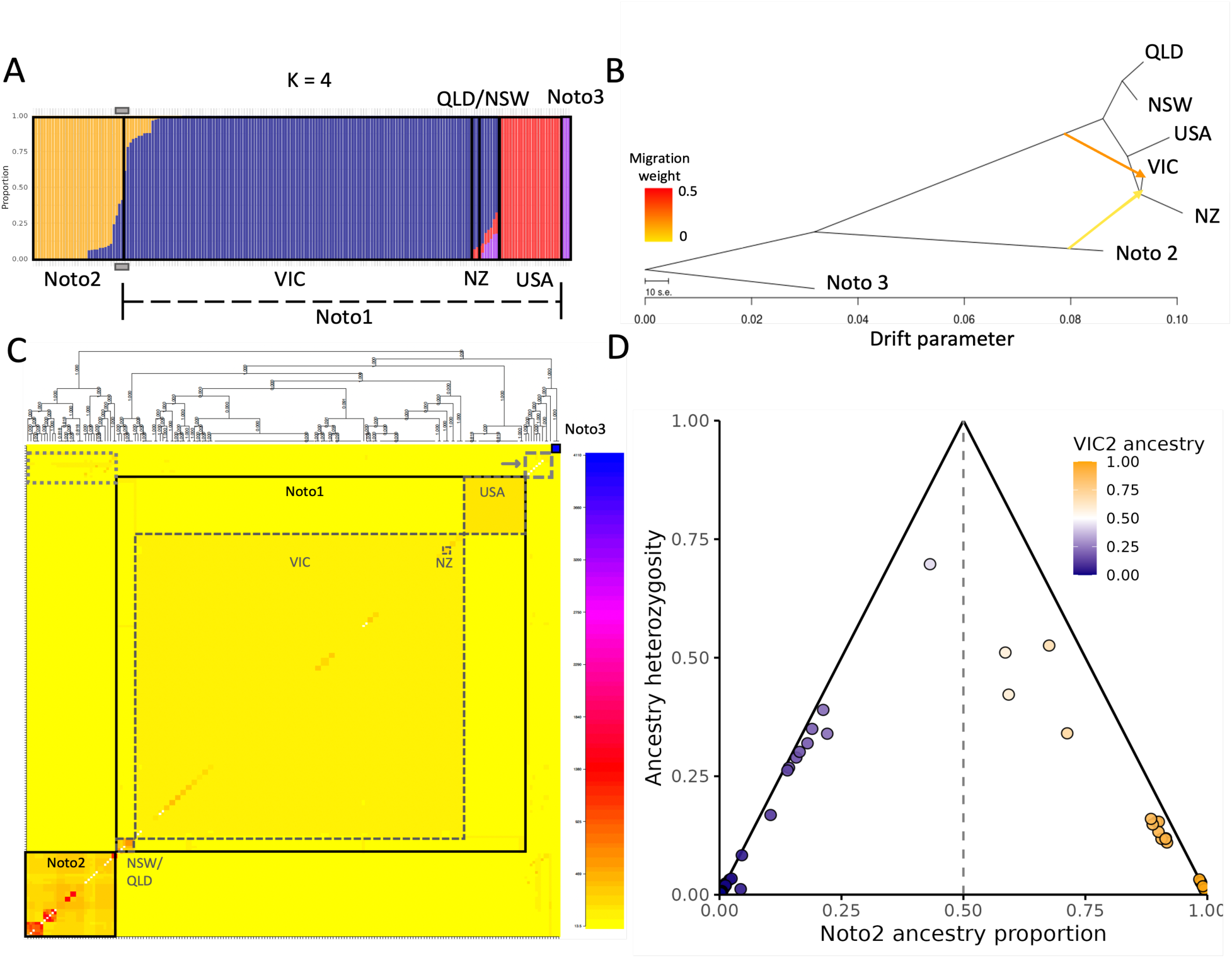
Population structure of Aedes notoscriptus across Australia, New Zealand, and the USA. (A) ADMIXTURE plot at K = 4 showing ancestry proportions for each individual, grouped by sampling region. (B) TreeMix analysis showing phylogenetic relationships and genetic drift among groups of Aedes notoscriptus. Arrows indicate migration edges, coloured by migration weight. (C) fineRADstructure co-ancestry heatmap and tree. Black solid boxes illustrate clustering of individuals into major groups (Noto1, Noto2, and Noto3), while grey dashed boxes show individual populations within Noto1. The light grey, long-dashed box with arrow indicates Noto1 individuals that do not cluster with other Noto1 samples. The light grey dotted box shows individuals with microhaplotypes between Noto1 and Noto2. (D) Triangle plot of ancestry heterozygosity versus Noto2 ancestry proportion for individual samples. Each point represents a single individual, with its position on the x-axis indicating the estimated proportion of ancestry derived from the Noto2 source population and its position on the y-axis indicating ancestry heterozygosity (the probability that alleles at a given locus derive from different ancestral sources). Solid black lines denote the theoretical maximum heterozygosity expected under a simple two-source admixture model, and the dashed vertical line marks equal (50:50) ancestry proportions. Points near the apex of the triangle represent individuals consistent with recent (e.g., F1) hybrids, whereas points near the base corners represent individuals with ancestry predominantly from one source population. Point colour reflects VIC2 (Noto2) ancestry proportion, from navy (low) through white (intermediate) to orange (high).

*Treemix* results also supported hybridisation between Noto1 and Noto2, including evidence of older admixture (Figure 3B). *Treemix* uses population allele frequencies to construct a maximum likelihood phylogeny that allows for migration edges to be placed between lineages within the phylogeny. Our analysis inferred two migration edges (Figure 3B). One of the two inferred migration edges linked Noto2 and Victorian Noto1 directly and might reflect recent hybridisation, while the other stronger migration edge was from deeper within the Victorian Noto1 clade and could reflect older admixture events or incomplete lineage sorting. We also used F3 tests to identify if any of the Victorian Noto1, Noto2, and Noto3 lineages were the product of admixture between the other lineages, finding no statistical evidence for this as all three populations had positive F3 values when tested with each other as sources (Victorian Noto1: f3 = 0.0411, Z = 35.99; Noto2: f3 = 0.0559, Z = 35.67; Noto3: f3 = 0.0678, Z = 45.82), though F3 tests can be underpowered with reduced-representation sequencing data as applied here (Malinsky et al. 2021).

Final evidence for admixture was through a *fineRADstructure* analysis, which analyses the pairwise similarity of RADtag microhaplotypes among all pairs of individuals (Figure 3C). In *fineRADstructure*, individuals are then ordered in x/y space by their similarity, while admixture shows up as microhaplotype similarity between individuals of different clusters that is displaced from the x/y diagonal (Figure 3C). As in other analyses of structure, *fineRADstructure* separated the main three lineages. However, a subset of 10 Victorian Noto1 individuals were not placed with other Noto1 individuals, instead occupying a space between the Noto1 and Noto3 clusters (Figure 3C, grey dashed box with arrow). Some of these individuals also had high microhaplotype similarity with Noto2 individuals (Figure 3C, grey dotted box). Interestingly, only one of the ten individuals included the putatively admixed individuals identified in *ADMIXTURE* and PCA (c.f. Figures 3A,S2), while the other nine all had >80% of Noto1 background and <20% Noto2 background as identified in *ADMIXTURE*.

These putative admixture patterns could reflect recent hybridisation, but could also reflect incomplete lineage sorting or uncertainty around assignment due to a lack of power (Linck and Battey 2019). To identify which of these explanations was more likely driving the observed patterns, we analysed levels of autosomal heterozygosity across individuals, where individuals that are the product of recent hybridisation should have higher heterozygosity. We considered that, for an individual to be most likely a recent (e.g. F1) hybrid, they should both appear admixed in one or more of the above analyses and have higher heterozygosity than unadmixed individuals.

Of the four putative hybrid samples identified through *ADMIXTURE* and PCA (Figure 3A, S2), three had much higher autosomal heterozygosity (0.0134, 0.0140, and 0.0188) than non-admixed samples from Victorian Noto1 (IQR = 0.00649—0.0077) and Noto2 (IQR = 0.00784—0.00101) (Figure 4A). The other putative hybrid had heterozygosity similar to Noto2 samples (0.0107). We consider these results to be evidence that the three highly heterozygous samples represent very recent hybridisation events and may be F1 hybrids of Victorian Noto1 and Noto2.

**Figure 4.**
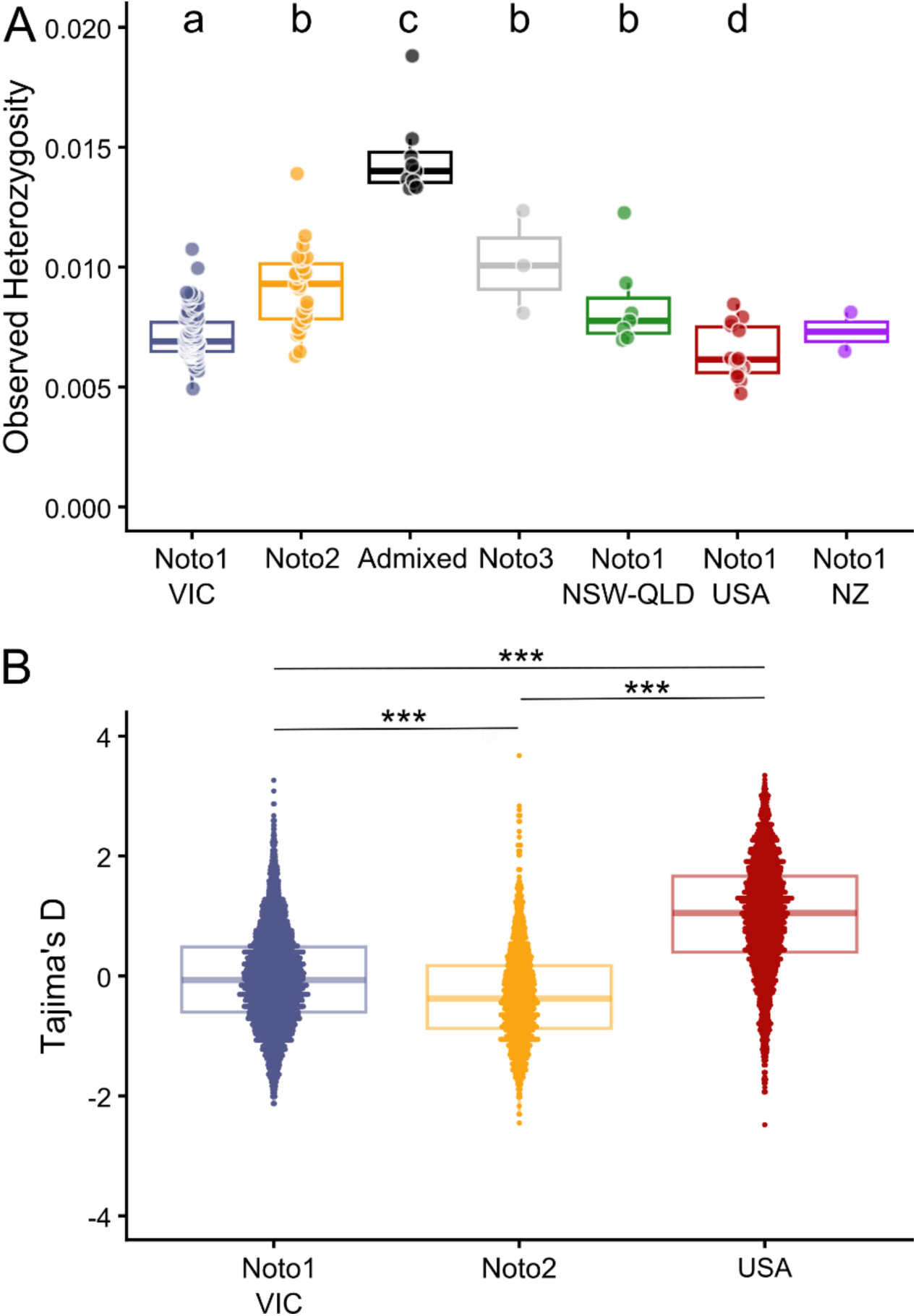
Genetic variation in Ae. notoscriptus. (A) Autosomal observed heterozygosity. Different letters (a-d) represent significant differences between groups (P < 0.05). NSW (n=4) and QLD (n=3) were combined to increase sample size. NZ had too few individuals (n=2) for statistical tests. (B) Tajima’s D in 100 kb windows, with populations rarified to n=13 and omitting windows with <5 SNPs. *** indicates highly significant differences (all P < 1 × 10^-16^; see main text).

We also investigated the 10 putative hybrids identified from *fineRADstructure* (Figure 3C). Nine of these had higher heterozygosity than non-admixed samples (range 0.0133 - 0.0188), while one had heterozygosity similar to Noto2 samples (0.0101). Notably, the putative hybrid identified by both *ADMIXTURE* and *fineRADstructure* had the highest heterozygosity of all the mosquitoes (0.0188). In line with our above criteria, we consider 11 individuals to be likely recent hybrids (nine identified through *fineRADstructure* and two through *ADMIXTURE*). A PCA of those 11 identified hybrids separated the nine samples identified from *fineRADstructure* away from the three samples identified through *ADMIXTURE* on PC1 (Figure S4). Note that we did not observe an obvious hybrid zone for Victorian Noto1 and Noto2 in the Greater Melbourne region, with hybrids occurring in the east, west and central part of the sampling area. However, five of the 11 admixed individuals were found at one single location in urban Melbourne (Figure 5C)

**Figure 5.**
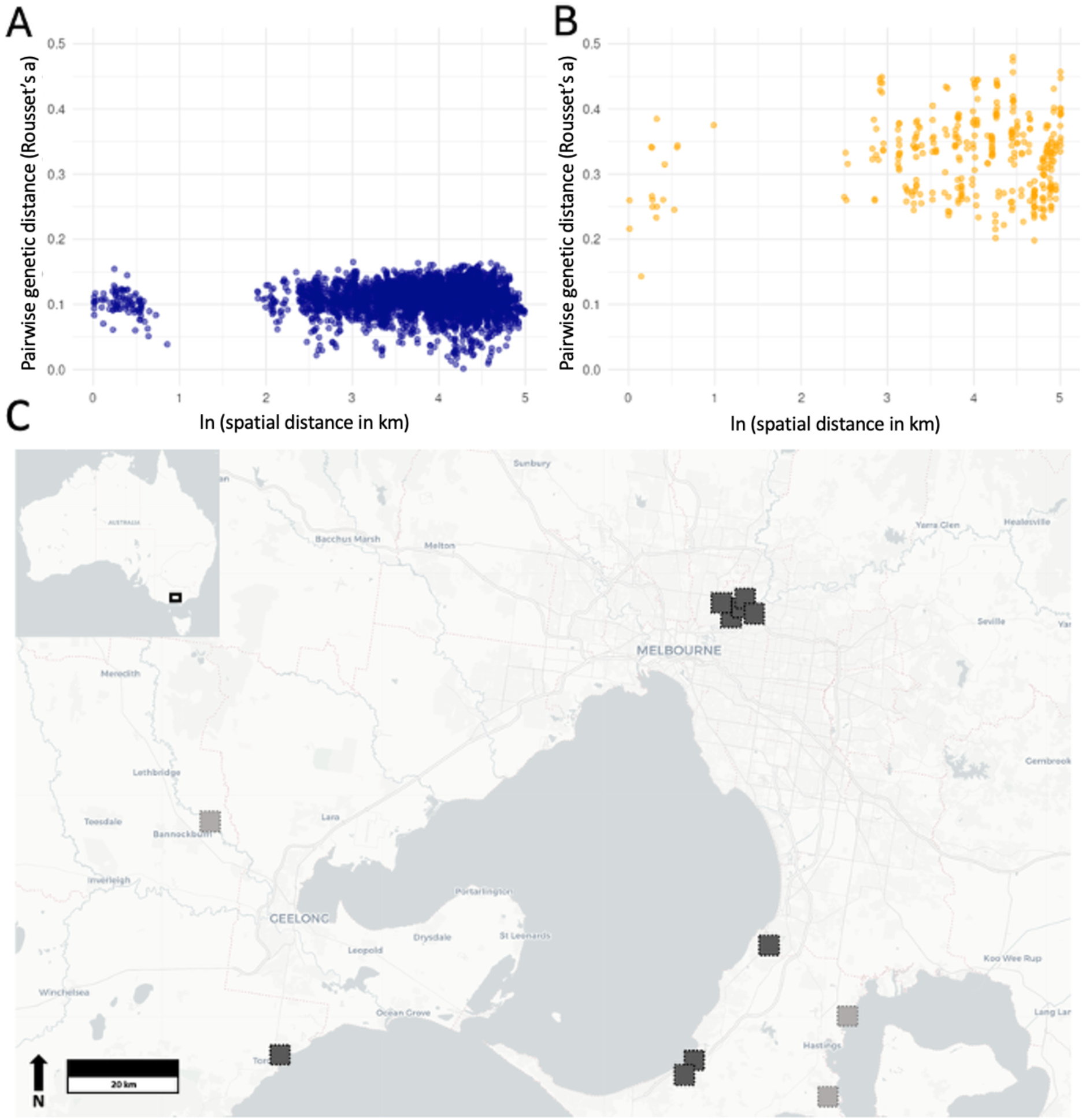
Scatter plot of a Mantel test between geographic (natural log) and genetic (Rousset’s a) distance for Victorian Noto1 (A) and Noto2 (B) without admixed individuals. (C) Locations of admixed individuals across the sampling area. Light grey squares show locations of admixed individuals identified in ADMIXTURE (Figure 3A+B) and dark grey squares show locations of individuals identified by fineRADstructure (Figure 3D).

Among the 43 sampling locations, Victorian Noto1 and Noto2 individuals co-occurred at 11 locations. Across these 11 locations, a total of 50 individuals were sampled (out of 155 individuals from all VIC locations), providing the subset of samples in which hybridisation was possible. Based on the observed local frequencies of Victorian Noto1 (*p*) and Noto2 (*q*) at these sites, the expected frequency of hybrids under random mating is 2*pq*, corresponding to an expected 41% hybrids, or approximately 20 individuals. In contrast, only 11 hybrid individuals were observed, representing 22% of individuals sampled at these locations. This deficit of hybrids relative to random mating expectations was statistically significant (one-sided exact binomial test: *p* = 0.005).

Site-specific expectations and observed counts are provided in Table S4. Because only a small number of individuals (≤5) were sampled per site, it is likely that additional locations also contained both Victorian Noto1 and Noto2 individuals but were not detected as such. Consequently, the contrast between the expected (≈20) and observed (11) number of hybrids may be conservative. Moreover, the absence of isolation by distance within either lineage across Greater Melbourne (section 2.6) and previously estimated strong dispersal in this species (Paris et al. 2023) suggests that movement above the scale of sampling sites may be possible, leading to additional opportunities for hybridisation. Accordingly, we repeated the analysis using the overall frequencies of Victorian Noto1 (120/155) and Noto2 (33/155) across the Greater Melbourne dataset, where the expected hybrid frequency under random mating (2*pq*) is approximately 33%, leading to ∼51 expected hybrids across Greater Melbourne. This deficit of hybrids relative to random mating expectations was highly statistically significant (one-sided exact binomial test: p = 1.8 × 10⁻¹⁵).

### 2.5 Evidence of a demographic bottleneck in the Victorian Noto1 lineage

We investigated autosomal observed heterozygosity (Schmidt et al. 2021A, Schmidt et al. 2024) within individuals and Tajima’s D (Tajima 1989) within populations, with Tajima’s D analysed in 100 kb windows and restricted to populations of sufficient sample size: Victorian Noto1, Noto2, and the USA (all rarified to n=13). Putative admixed individuals were analysed separately. Autosomal heterozygosity was used as a metric of genome-wide genetic variation and is expected to decrease when populations experience demographic bottlenecks. Tajima’s D summarises the shape of the folded site frequency spectrum, where increasingly positive scores indicate a greater proportion of common alleles than expected under demographic equilibrium, as may occur when populations experience demographic bottlenecks.

There were clear and statistically significant differences in heterozygosity among Australian lineages and between Australian lineages and the invasive USA lineage (Figure 4A). Among the Australian lineages, heterozygosity was lowest in Noto1 (Victoria, x̄ = 0.00716; NSW and QLD, x̄ = 0.00841). Heterozygosity in Victorian Noto1 was 21.5% lower than Noto2 (x̄ = 0.00913 Student’s t-test, t_116_ = 7.78, P = 3.3 × 10^-12^), 29.8% lower than Noto3 (x̄ = 0.0102; Student’s t-test, t_92_ = 5.11, P = 1.8 × 10^-6^), and 14.8% lower than NSW and QLD Noto1 (Student’s t-test, t_96_ = 3.05, P = 0.003) (Figure 2B). Heterozygosity differences were less pronounced between Victorian Noto1 and the recently established, invasive USA population (x̄ = 0.00642; 10.3% lower; Student’s t-test, t_104_ = 2.69, P = 0.008). As the low heterozygosity in USA presumably reflects a genetic bottleneck from this invasion, these results point to another possible bottleneck in the Noto1 lineage that would explain its lower heterozygosity relative to the Noto2 and Noto3 lineages (Figure 4A). By comparison, a ∼35% reduction in heterozygosity was recorded between native and invasive *Ae. aegypti* populations (Kent et al. 2025), a difference equivalent to that between USA and the Noto3 population (36.9%).

Patterns of Tajima’s D were also in support of a demographic bottleneck in Victorian Noto1 (Figure 4B). The contraction phase of a bottleneck will lead to elevated Tajima’s D values and increased variance relative to populations at demographic equilibrium (Hahn 2018). As expected, the recently established invasive USA population had higher mean and variance in Tajima’s D than Victorian Noto1 (USA: x̄ = 1.000, σ^2^ = 0.867; Noto1: x̄ = -0.031, σ^2^ = 0.643; Student’s t-test, t_7094_ = 50.0, P < 1 × 10^-16^). However, Victorian Noto1 had higher mean and variance than Noto2 (Noto2: x̄ = -0.320, σ^2^ = 0.604; Student’s t-test, t_5972_ = 13.3, P < 1 × 10^-16^). Genomic estimates of Tajima’s D in mosquitoes are typically negative in native range populations (Tennessen et al. 2021, Love et al. 2023, Boddé et al. 2025, Schmidt et al. 2025), potentially reflecting global mosquito population expansions.

Taken together, the combination of reduced heterozygosity and elevated Tajima’s D in Victorian Noto1 relative to Noto2 supports the inference that Victorian Noto1 has experienced one or more demographic bottlenecks. One bottleneck may have occurred in the broader Noto1 lineage, potentially coinciding with divergence from Noto2, followed by a more recent bottleneck specific to Victoria that may reflect a range expansion into the Greater Melbourne region. In contrast, Noto2 shows no evidence of reduced genetic diversity or distortion of the site-frequency spectrum, consistent with the Greater Melbourne region representing part of its ancestral range.

### 2.6 No isolation by distance across Greater Melbourne

Isolation by distance across the Greater Melbourne region was tested separately for the Victorian Noto1 and Noto2 lineages after putative admixed individuals were removed. Mantel tests comparing pairwise genetic distances (Rousset’s α) with the natural logarithm of geographic distances revealed no significant relationship for either Victorian Noto1 (Mantel test: *r* = 0.0219, *p* = 0.0752) or Noto2 (Mantel test: *r* = 0.0128, *p* = 0.3671), indicating an absence of detectable isolation by distance within either lineage at this spatial scale (Figure 5A,B). The lack of isolation by distance within both lineages could result from high dispersal in both lineages across the Greater Melbourne region, resulting in near-panmixia at the scales sampled.

However, this result could also be influenced by inter-lineage introgression of divergent genetic backgrounds through hybridisation which, though clearly restricted by reproductive isolation, is nevertheless occurring across Greater Melbourne and may interfere with isolation by distance patterns. Hybridisation between Victorian Noto1 and Noto2 is geographically widespread (Figure 5C), and although recently admixed individuals were excluded from the isolation-by-distance analyses, the effects of admixture in previous generations may still affect these patterns. This admixture could in part explain the high pairwise genetic distances among Noto2 individuals even over short distances (Figure 5B).

### 2.7 A single massive structural variant segregating in one Victorian lineage

#### 2.7.1 Identification of structural variants from genome-wide outlier windows

To identify structural variation such as inversions, we used a window-based local PCA method run in *lostruct* (Li and Ralph 2019). Local PCA identifies regions of the genome containing clusters of MDS outliers, and these outlier regions can be specifically investigated for signs of structural variation. Our analysis found a single large continuous outlier region on chromosome 2 within the Noto2 group (Figure 6A). A subsequent PCA of the 5200 SNPs within this outlier region revealed three discrete clusters of individuals (Figure 6B). To assess whether the elevated signal in this region could be driven by variation in data completeness, we compared per-window missingness between the putative inversion region and the background of chromosome 2. Mean window-based missingness was only slightly higher within the inversion region (0.589) compared to background genomic regions (0.491). This small difference was unlikely to explain the strong population structure observed within the region.

**Figure 6.**
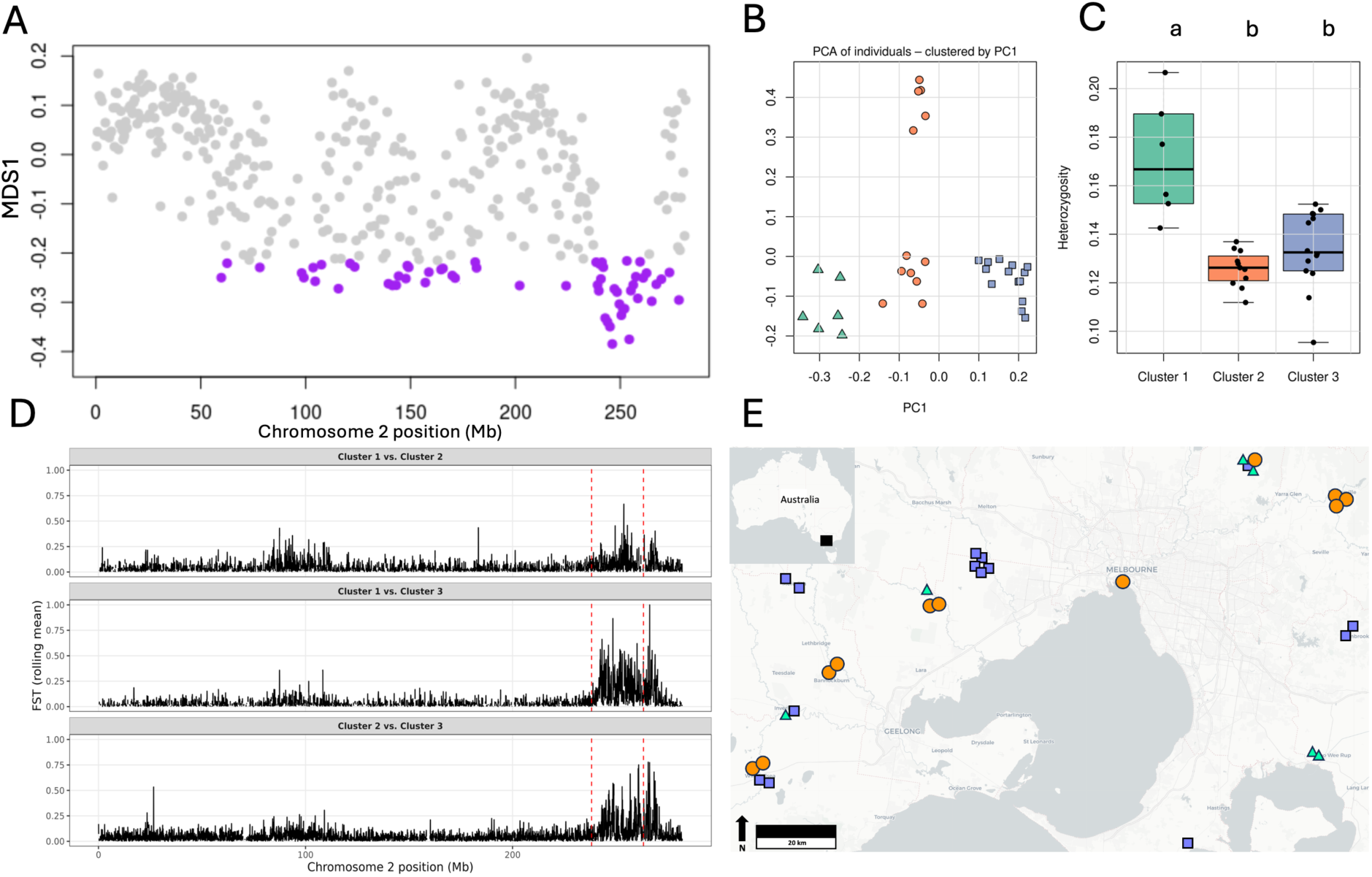
Characterisation of the MDS outlier region on chromosome 2 of Noto2 lineage. (A) Plot of MDS values across chromosome 2. Each grey dot represents a window of 50 SNPs and outlier windows are highlighted in red. (B) PCA based on SNPs from outlier region. Three clusters identified and illustrated in green triangles, orange circles and blue squares. (C) Heterozygosity for each of the groups identified in PCA. (D) The rolling average of pairwise F_ST_ between the three clusters across chromosome 2. (E) Map indicating the locations of the clusters identified in PCA.

If this region represents an inversion, we expect the two clusters to represent the two homokaryotypes of the inversion, with the third cluster representing the heterokaryotypic individuals. Individual heterozygosity differed significantly among PCA-defined clusters (one-way ANOVA: F₂,₂₈ = 16.71, p = 1.7 × 10⁻⁵). Post hoc Tukey tests showed that cluster 1 exhibited significantly higher heterozygosity than both cluster 2 (p = 1.5 × 10⁻⁵) and cluster 3 (p = 1.4 × 10⁻⁴), whereas clusters 2 and 3 did not differ significantly from each other (p = 0.43). This pattern is consistent with cluster 1 representing heterokaryotypes and clusters 2 and 3 representing alternative homokaryotypes (Figure 6C) (Huang et al. 2020).

Pairwise *F_ST_* comparisons among the three local PCA clusters revealed strong differentiation in the outlier region on chromosome 2. The greatest divergence was observed between the putative heterokaryotypic cluster 1 and the homokaryotypic cluster 3 (mean *F_ST_* = 0.32), followed by differentiation between the two putative homokaryotypes, clusters 2 and 3 (mean *F_ST_* = 0.24; Figure 6D). In contrast, differentiation between the heterokaryotypic cluster 1 and homokaryotypic cluster 2 was more moderate (mean *F_ST_* = 0.16). Empirical permutation tests comparing putative inversion-region *F_ST_* to randomly sampled genomic regions of equal size across chromosome 2 indicated that differentiation in the outlier region was significantly elevated above the chromosomal background for cluster 1 vs. cluster 3 and cluster 2 vs. cluster 3 (p < 0.001 in both cases), whereas the increase for cluster 1 vs. cluster 2 was weaker and not statistically significant (p = 0.075; Figure 6D).

We next evaluated whether the highly differentiated genomic region consistent with a putative inversion exhibited patterns of spatial or lineage-specific structure. Although inversions in mosquitoes can show geographic clines or localised differentiation (Ayala et al. 2017, Cheng et al. 2018), we observed no clear spatial structuring of putative karyotypes across Greater Melbourne (∼200 km), with both homokaryotypes occurring across eastern, western, northern, and southern localities (Figure 6E). This suggests that, at this spatial scale, the structural variant is not geographically partitioned.

Because the putative inversion was observed segregating only in Noto2, we further examined differentiation between each Noto2 cluster and Victorian Noto1 across the outlier region. Pairwise comparisons showed that cluster 2 exhibited consistently low differentiation relative to Noto1 (mean *F_ST_* = 0.037) and no significant elevation above genomic background (permutation test, p = 0.12). In contrast, clusters 1 and 3 showed significantly elevated differentiation within the outlier region compared to Victorian Noto1 (mean *F_ST_* = 0.050 and 0.069, respectively; permutation tests, p < 0.001). These results indicate that Victorian Noto1 most closely matches the putative inversion orientation present in Noto2 cluster 2, whereas Noto2 cluster 3 likely represents the alternative arrangement and cluster 1 the heterokaryotype. The elevated differentiation confined to this region, relative to background levels, is consistent with recombination suppression maintaining distinct haplotypic classes rather than genome-wide divergence.

We were also curious whether we missed MDS outlier regions if they were driven by variants present at low frequency and therefore repeated the local PCA on the Victorian Noto1 and Noto2 dataset without applying the −R filtering step in *Populations*. Under this relaxed filtering, we identified two additional outlier regions on chromosome 1 in Noto2 (Figure S5). However, these signals were largely driven by individuals from a single sampling location, and the corresponding PCA patterns did not show clear or consistent clustering (Figure S5). Given the small number of individuals contributing to these outliers and the absence of supporting evidence from *F_ST_* or heterozygosity analyses, we cannot determine whether these regions represent biologically meaningful signals or artefacts of sampling and low allele counts, and we therefore treat them with caution.

#### 2.7.2 Functional enrichment of neuronal signalling and oxidoreductase genes within the chromosome 2 outlier region

BLAST analysis of sequences from the outlier region against the *Ae. aegypti* reference genome revealed approximately 25 Mb of contiguous sequence overlapping multiple annotated genes. To summarise gene content and estimate the number of individual genes within this region, we counted the number of BLAST hits per gene or transcript. The top hits corresponded to genes with multiple transcript variants, including *attractin*, *serine/threonine-protein kinase 32A*, *calpain-D*, and *tyrosine-protein kinase receptor torso* (Table S5). The number of BLAST hits per gene varied from 1 to >4,600, reflecting multiple transcript isoforms and high sequence coverage. Using the median gene length across the region, we estimate that the outlier region contains approximately 112 individual genes.

To investigate the potential functional roles of candidate genes within the chromosome 2 outlier region, we performed Gene Ontology (GO) enrichment analysis using g:Profiler. Genes located within the outlier region were used as the query set, and enrichment was assessed relative to the full *Ae. aegypti* genome annotation. Significance was determined using g:Profiler’s built-in multiple-testing correction (g:SCS), with adjusted p < 0.05 considered significant (Table 1). Seven molecular function categories met this threshold, several of which were hierarchically related and therefore not statistically independent.

**Table 1.** Significantly enriched molecular functions and pathways for candidate genes in the chromosome 2 outlier region (g:Profiler Analysis)

| CATEGORY | TERM NAME | TERM ID | ADJUSTED P-VALUE | −LOG10 |
| --- | --- | --- | --- | --- |
| GO:MF | Transmitter-gated monoatomic ion channel activity | GO:0022824 | $2.06 \times 10^{-32}$ | 31.99 |
| GO:MF | Transmitter-gated channel activity | GO:0022835 | $2.06 \times 10^{-32}$ | 31.99 |
| GO:MF | Extracellular ligand-gated monoatomic ion channel activity | GO:0005230 | $9.76 \times 10^{-32}$ | 31.01 |
| GO:MF | 2-alkenal reductase [NAD(P)+] activity | GO:0032440 | $1.28 \times 10^{-21}$ | 20.89 |
| GO:MF | 15-oxoprostaglandin 13-oxidase [NAD(P)+] activity | GO:0047522 | $1.28 \times 10^{-21}$ | 20.89 |
| GO:MF | Neurotransmitter receptor activity | GO:0030594 | $1.49 \times 10^{-21}$ | 20.83 |
| GO:MF | Oxidoreductase activity, acting on CH-CH group of donors, NAD or NADP as acceptor | GO:0016628 | $2.72 \times 10^{-21}$ | 20.57 |
| KEGG | Polycomb repressive complex | KEGG:03083 | $2.79 \times 10^{-21}$ | 20.55 |

Enriched neuronal signalling functions included ligand-gated ion channel activity and neurotransmitter receptor activity, corresponding to Cys-loop receptor families such as nicotinic acetylcholine and GABA receptors that mediate rapid synaptic transmission (Matthews et al. 2016, Jones 2018), and are implicated in sensory-driven behaviours including host-seeking in *Ae. aegypti* (Matthews et al. 2016). Oxidoreductase-related terms were also enriched, including 2-alkenal reductase and 15-oxoprostaglandin oxidoreductase activities, consistent with redox processes that buffer oxidative stress during blood feeding and influence fitness-related traits in *Aedes* mosquitoes (Oliveira et al. 2011, 2017, Bottino-Rojas et al. 2018). KEGG pathway analysis further indicated enrichment of components of the Polycomb repressive complex, a chromatin regulatory system associated with H3K27me3-mediated transcriptional control in mosquitoes (Vilcinskas 2017).

## 3 Discussion

Detecting and characterising reproductive isolation have seen a resurgence of interest in the current genomic era (Feder et al. 2012, Westram et al. 2022, Yoshida et al. 2023, van der Heijden et al. 2025). Here, we investigated patterns of reproductive isolation in the major international mosquito pest *Ae. notoscriptus*, a species of growing public health concern following its implication in the transmission of Buruli ulcer in Australia (Mee et al. 2024) and its recent invasion and spread throughout California (Metzger et al. 2021). Using genome-wide reduced representation sequencing across both the native and invasive range, together with a chromosome level reference genome, we identified three deeply divergent native lineages, including two that coexist sympatrically across Greater Melbourne. Despite extensive geographical overlap, these lineages exhibited incomplete reproductive isolation, with fewer hybrids observed than expected under random mating.

The mito-nuclear discordance observed in *Ae. notoscriptus* (Figure 2B) has likely been produced by hybridisation followed by backcrossing between lineages. Although incomplete lineage sorting can generate similar patterns in large populations and may have contributed to patterns here, the widespread sympatry of divergent lineages and the presence of admixed individuals make mitochondrial introgression via hybridisation a likely feature in this system (Palumbi et al. 2001, Toews and Brelsford 2012). Even rare hybridisation events can facilitate the transfer of mtDNA across lineage boundaries, decoupling mitochondrial and nuclear ancestry despite overall selection against hybrids. Regardless of the precise mechanism, this discordance highlights the limitations of mtDNA for delineating lineage boundaries in *Ae. notoscriptus* and exemplifies the need for genome-wide or targeted nuclear markers for accurate epidemiological surveillance.

Turning to the nuclear lineages, these divergent genomic backgrounds differed strikingly in their population genetic characteristics. The more widespread Noto1 lineage showed reduced autosomal heterozygosity and elevated Tajima’s D relative to Noto2 (Figure 4), consistent with one or more past demographic bottlenecks. In contrast, Noto2 retained higher genetic diversity indicative of longer-term persistence in the region. These contrasting signatures suggest that the Greater Melbourne region likely represents part of the ancestral range of Noto2, while Noto1 may have expanded into the region more recently following population contraction. The sympatric occurrence of Victorian Noto1 and Noto2 across Greater Melbourne raises the question of how these divergent lineages arose and persist despite ongoing contact. Despite extensive geographic overlap, hybrids were observed at significantly lower frequencies than expected under random mating, indicating partial but incomplete reproductive isolation. Such isolation is likely maintained by a combination of pre-mating barriers, including divergence in mating behaviour through activity timing (Sawadogo et al. 2013), host preference (Haba and McBride 2022), or microhabitat use (Diabaté et al. 2009, Gimonneau et al. 2012), as well as post-mating barriers such as reduced hybrid fitness (Deitz et al. 2020, Zheng 2020). The presence of large-scale genomic differentiation between lineages suggests that multiple isolating mechanisms act together to limit gene flow in sympatry. While no obvious morphological differences among lineages were observed during processing of samples, this study did not investigate these carefully, and morphological differences should be considered in future research.

Although we found evidence of reproductive isolation, it was not absolute, and even rare hybridisation events may permit the transfer of alleles between lineages, particularly if favoured by selection. Despite possible ongoing selection against hybrids, such introgression could facilitate the spread of traits relevant to pest management, including insecticide resistance (Norris et al. 2015), altered vector competence (Fontaine et al. 2015), or adaptation to new climate conditions (Cheng et al. 2012). *Wolbachia*-induced cytoplasmic incompatibility can be excluded as a mechanism contributing to reproductive isolation in *Ae. notoscriptus*, as no individuals in our dataset carried these infections. This is consistent with previous studies in Victorian populations (Bellarine Peninsula and Melbourne), which showed extremely low *Wolbachia* prevalence (Duchemin et al. 2017), in contrast to a higher prevalence reported in some Queensland populations (Skelton et al. 2016). The broader geographic prevalence of *Wolbachia* in this species warrants future investigation.

Recent work on *Ae. notoscriptus* challenged the view that this species is primarily locally structured by reporting evidence for long-range dispersal and weak spatial genetic structure across a Melbourne area (Paris et al. 2023). In that study, isolation by distance was no longer detected once closely related individuals were removed, consistent with high effective dispersal and near-panmixia at the spatial scale sampled. The results presented here are consistent with this interpretation. When analysed separately, neither Victorian Noto1 nor Noto2 exhibited significant isolation by distance across Greater Melbourne after putative admixed individuals were removed (Figure 5), which may reflect high dispersal rates across Greater Melbourne. Alternatively, the lack of isolation by distance may reflect the latent admixture among the lineages resulting from occasional hybridisation. The key additional insight from this study is that such spatial panmixia can coexist with reproductive isolation between sympatric lineages. While Victorian Noto1 and Noto2 are not spatially structured within the region, they nonetheless remain genetically distinct due to barriers to gene flow that are not primarily geographic in nature. High levels of effective dispersal in *Ae. notoscriptus* may reflect not only flight capacity but also ecological or behavioural traits that increase association with human-modified environments (Kay et al. 1979, Takken and Verhulst 2013), and whether such traits differ between Noto1 and Noto2 remains an important open question. The widespread geographic distribution of admixed individuals (Figure 5C) confirms that Greater Melbourne represents an active zone of secondary contact.

Chromosomal inversions are increasingly recognised as important features of mosquito genomes and have been linked to ecological differentiation, including associations with climate, latitude, and adaptive traits across multiple species complexes (e.g. Ayala et al. 2011, 2014, 2017, Fouet et al. 2012, Liang et al. 2025). In *Ae. notoscriptus*, we identified a massive structural variant spanning approximately 25 Mb (Figure 6) and containing an estimated ∼120 genes enriched for neuronal signalling, redox and metabolic processes, and chromatin-mediated regulation (Table 1); pathways directly associated with host-seeking behaviour (Matthews et al. 2016), activity timing (Sawadogo et al. 2013), reproductive output, and physiological stress responses (Diabaté et al. 2009, Gimonneau et al. 2012).

Chromosomal rearrangements such as inversions can contribute to reduced hybrid frequencies through suppressed recombination and the accumulation of incompatibilities in rearranged regions, potentially leading to reduced fertility or viability of heterokaryotypes (e.g. Ayala et al. 2013, 2014; Dickson et al. 2016; Feder and Nosil 2009; Lowry and Willis 2010). However, investigating how this may be contributing to the isolation of Noto1 and Noto2 is beyond this present study. Confirming whether this variant represents an inversion will require whole-genome sequencing to resolve breakpoint structure and cytogenetic validation to determine orientation directly (George et al. 2010). Broader geographic sampling including of the Noto3 lineage will be essential to clarify the variant’s evolutionary origin, reconstruct whether the derived arrangement arose within Noto2 or is ancestral and fixed in Noto1, and distinguish true parental genotypes from later-generation backcrosses that cannot be reliably separated using reduced-representation data alone.

The positionally-informed genomic analyses presented here have been enabled by our chromosome-level reference assembly. *Aedes* genomes are notoriously difficult to assemble owing to high repeat content and are ∼5 times larger than those of *Anopheles* mosquitoes; there are thus far fewer genomes assembled for *Aedes* species than *Anopheles* (Schmidt et al. 2021B). Our assembly for *Ae. notoscriptus* is one of ∼5 high-quality genomes available for *Aedes* globally (Dudchenko et al. 2017, Deng et al. 2024, Morinaga et al. 2025) and is only the second chromosome-level assembly for an endemic Australian mosquito after *An. farauti* (Waterhouse et al. 2020). Beyond enabling the analyses presented here, this resource provides a foundation for future comparative genomic work across *Aedes* that may shed light on phenotypic differences including vectorial capacity, as will future genome assemblies produced for the other *Ae. notoscriptus* lineages identified in this cryptic complex (Noto2 and Noto3). More broadly, expanded geographic sampling across both the native and invasive ranges will be important to determine whether additional cryptic lineages exist and to resolve the precise origins and invasion pathways of international populations.

## Conclusion

Our study reveals that *Ae. notoscriptus* comprises deeply divergent cryptic lineages that can coexist in sympatry while remaining significantly, though incompletely, reproductively isolated. By integrating genome-wide data with a chromosome-level reference assembly, we show that differences in demographic history, dispersal dynamics, and large-scale genomic architecture likely interact to shape lineage boundaries in this species. These findings highlight the limitations of mitochondrial markers for lineage identification in this species and emphasise the need for nuclear genomic tools in epidemiological surveillance and vector management. More broadly, our results show cryptic evolutionary diversity within mosquito pests which may influence dispersal, control efficacy, and disease risk, reinforcing the importance of incorporating evolutionary genomics into applied public health strategies.

## 4. Material and Methods

### 4.1 Reference genome assembly

#### 4.1.1 DNA extraction and sequencing

We selected a single *Ae. notoscriptus* male adult from Victoria, Australia, later confirmed as of the Noto1 lineage (see Results). High molecular weight genomic DNA was extracted using the Nanobind Tissue Big DNA Kit (Cat#: NB-900-701-01, Circulomics) and the High Molecular Weight Insect DNA Extraction Protocol. The DNA sample was then submitted to the Biomolecular Resource Facility (John Curtin School of Medical Research, Australian National University) for long read sequencing on the PacBio Revio platform. Briefly, the genomic DNA was sheared using the Megaruptor 3.0 (Diagenode B06010003), then size-selected to deplete for DNA fragments less than 8 kb using the Sage Science BluePippin (cassette kit: PAC20KB, gel: 0.75% agarose, marker: S1). The genomic DNA was then amplified using the SMRTbell gDNA sample amplification kit (Pacific Biosciences, Cat #: 101-980-000) and used to prepare a SMRTbell library using the SMRTbell Prep Kit 3.0 (Pacific Biosciences, Cat: PCB-102-182-700). The SMRTbell library was then sequenced on a single SMRT cell (Pacific Biosciences, Cat: PCB-102-202-200).

A pool of two frozen *Ae. notoscriptus* adults from the Noto1 lineage was submitted to the Biomolecular Resource Facility (John Curtin School of Medical Research, Australian National University) for chromatin conformation capture using the Arima Hi-C Kit for animal tissues (Arima Genomics, CA, USA, Cat# A510008) according to the manufacturer’s instructions.

Briefly, tissue was crosslinked and digested with a restriction enzyme cocktail. The 5’ overhangs were filled in and labelled with biotinylated nucleotides. Spatially proximal DNA were ligated and purified. The proximally-ligated DNA was then fragmented and enriched for biotinylated fragments. The Illumina library was prepared using the Kapa Hyper Prep kit (ARIMA Document Part Number A160139 v00) and sequenced on an S4 flow cell of an Illumina NovaSeq6000 in paired-end mode (2x 150bp PE).

#### 4.1.2 Genome assembly

The genome was assembled using mabs-hifiasm.py in *Mabs* v2.2 (Schelkunov 2023) with the Diptera odb10 BUSCO lineage, integrating HiFi and Hi-C data under default parameters.

Redundant haplotigs were removed using *Purge_Dups* v1.2.5 (Guan et al. 2020) with HiFi reads, and the assembly was decontaminated using the *NCBI Foreign Contamination Screening* v0.5.0 (FCS) pipeline (Astashyn et al. 2024) using taxid 508665. Chromosome-scale scaffolding was performed with *ALLHiC* v0.9.14 (Zhang et al. 2019) using Hi-C data, followed by manual curation in *Juicebox* v2.17.0 (Durand et al. 2016). The final assembly is available in GenBank under accession GCA_040801935.1.

Genome completeness was assessed using *BUSCO* v5.2.2 (Benchmarking Universal Single-Copy Orthologs) (Simão et al. 2015, Manni et al. 2021) against the *Diptera odb10* and *Insecta odb10* lineage datasets (Kriventseva et al. 2019). Analyses were performed in genome mode using default parameters.

### 4.2 Sampling and DNA extraction

We obtained samples of *Ae. notoscriptus* from 58 locations from Australia: Victoria (VIC), New South Wales (NSW), the Australian Capital Territory (ACT), Queensland (QLD) and the Northern Territory (NT) as well as from New Zealand (NZ) and from the United States of America (USA) (see Table S6; Figure 1). Most collections deployed ovitraps that consisted of a black 1.2L plastic bucket, halfway filled with water containing several alfalfa pellets to attract gravid *Ae. notoscriptus* (Ritchie 2001). A strip of red felt extending into the water provided an oviposition substrate. Felts were collected after 7 days and partially dried.

Three days after collection, eggs were hatched in 500 mL reverse osmosis (RO) water containing 2-3 TetraMin® tropical fish food tablets (Tetra, Melle, Germany). Water and food were replaced as appropriate. Samples from NSW, ACT, QLD, NZ, and USA were reared from larvae collected directly from the environment and *Ae. notoscriptus* from the NT were collected as adults using BG traps (Biogents AG, Regensburg, Germany). Adults were transferred into absolute ethanol and stored at -20°C until DNA extraction.

Keys from Dobrotworsky (1965) were used to identify *Ae. notoscriptus* mosquitoes using morphological features. DNA was extracted from a subset of individual mosquitoes per location (Table S6) using either Qiagen DNeasy Blood & Tissue Kits (Qiagen, Hilden, Germany) or Roche High Pure PCR Template Preparation Kits (Roche Molecular Systems, Inc., Pleasanton, CA, USA).

### 4.3 DNA sequencing and library construction

We created double-digest restriction site-associated DNA sequencing (ddRADseq) libraries following the method for *Ae. aegypti* as described by Rašić et al. (2014). We began by digesting 10 ng of genomic DNA with 10 units each of MluCI and NlaIII restriction enzymes, using NEB CutSmart buffer (New England Biolabs) and water. The digestions were conducted at 37°C for 3 hours without a heat kill step and the products were cleaned using paramagnetic beads (Sera-Mag Magnetic Carboxylate modified microparticle solutions, GE Healthcare, Chicago, USA). Modified Illumina P1 and P2 adapters were then ligated onto the cleaned digestions overnight at 16°C with 1000 units of T4 ligase (New England Biolabs), followed by a 10-minute heat deactivation step at 65°C. We performed size selection using a Pippin-Prep 2% gel cassette (Sage Sciences) to retain DNA fragments of 350-450 bp.

The size-selected libraries were amplified by PCR, using 1 μL of the size-selected DNA, 5 μL of Phusion High Fidelity 2× Master mix (New England Biolabs) and 2 μL of 10 μM standard Illumina P1 and P2 primers. The PCR reactions were conducted for 12 cycles, then cleaned and concentrated using 0.8x paramagnetic beads. Each ddRAD library contained 24 mosquitoes and each was sequenced on a single sequencing lane using 150 bp chemistry. Libraries were sequenced 150 bp paired-end at GeneWiz Inc (Suzhou, China) using a HiSeq 4000 (Illumina, California, USA).

### 4.4 Genotyping and filtering of ddRADseq data

#### 4.4.1 Data processing and read alignment

We used the *Process_radtags* program in *Stacks* v2.0 (Catchen et al. 2013) to separate the sequence reads by sample. Reads with low quality were discarded using a 15 bp sliding window if the average Phred score fell below 20. We then used *Bowtie* v2.0 (Langmead and Salzberg 2012) to align reads the reference genome assembled from a Noto1 individual (as described in section 4.1) using the "very-sensitive" alignment settings. We filtered the resulting bam files to include only paired reads that aligned concordantly, requiring the two paired reads to align to the same contig to avoid multi-mapping using *Samtools* (Danecek et al. 2021). These above protocols are in line with current understandings around limiting reference bias (Akopyan et al. 2025).

#### 4.4.2 Genotyping and filtering for genetic structure and local PCA

Stacks *Ref_map* program was used to build *Stacks* catalogs, from which genotypes were called at a 0.05 significance level and a minimum mapping quality of 15 to filter any remaining multi-mapped reads. We generated VCF files for the catalog with the *Stacks* program *Populations* (Catchen et al. 2013). Multiple datasets were generated for different analyses:

• **Global dataset:** All individuals included, no prior population assignment, retaining loci present in at least 85% (-R 0.85) of individuals and with a minimum allele count of 3 (--min-mac 3), yielding 227,982 SNPs across 189 individuals, used for PCA (section 4.5.1) and *ADMIXTURE* (section 4.5.2) and *fineRADstructure* (section 4.5.3).

• **Seven populations dataset:** Populations defined based on initial clustering, with an additional minimum 25% locus presence per population (-p 7; -r 0.25), retaining 226,298 SNPs from 189 individuals, used for *Treemix* (section 4.5.4) analyses.

• **Australian-only dataset:** Excluding invasive populations from the USA and NZ to focus on population splits in the native range, retaining 266,659 SNPs from 166 individuals, used for DAPC (section 4.5.1) analysis.

• **Victorian Noto1 and Noto2 datasets:** Analyses restricted to these populations for local population structure and isolation-by-distance (IBD) analyses (section 4.5.5) and the detection of structural variants (section 4.5.8), retaining 181,966 SNPs from 122 Noto1 individuals and 202,787 SNPs from 33 Noto2 individuals.

Missingness was generally low across the dataset and broadly consistent with expectations for ddRAD sequencing. Noto1, which was also used as the reference lineage for read mapping, showed the lowest missingness and highest mean sequencing depth. Higher missingness was observed in more divergent lineages, including Noto2 and Noto3 (Table S1) Across most populations, missingness remained low and did not show strong separation among groups. Sequencing depth varied among populations but remained within a similar order of magnitude across the dataset.

#### 4.4.3 Genotyping and filtering for heterozygosity, Tajima’s D and phylogenetics

The following pipeline was used for analysis of autosomal heterozygosity (section 4.5.7), Tajima’s D (section 4.5.7) and d*_XY_* (section 4.5.6). Here it was of critical importance to obtain accurate genotypes for monomorphic sites and rare alleles including singletons (Schmidt et al. 2021A). We used a published pipeline for autosomal heterozygosity (Schmidt et al. 2024) to genotype filtered bam files from ***4.4.2*** and process them further. This pipeline genotypes all individuals together, applies hard filters to all individuals together, and then exports VCF files for each individual or subset of individuals for further filtering by sequencing depth, missing data, star alleles and spanning deletions. GVCF files were produced for each individual using HaplotypeCaller in GATK v4.2.6.1 (Van der Auwera & O’Connor, 2020), and these were genotyped using GenotypeGVCFs set to ‘--include-non-variant-sites’. Hard filtering excluded indels then followed standard GATK guidelines for non-model taxa, setting ‘QD < 2.0’, ‘QUAL < 30.0’, ‘SOR > 3.0’, ‘FS > 60.0’ and ‘MQ < 40.0’.

### 4.5 Population structure

#### 4.5.1. PCA and DAPC

We used *PLINK* v. 1.90b6.16 (Purcell et al. 2007) to convert the VCF genotype file of the global dataset to PLINK binary format. Principal component analysis (PCA) was performed in PLINK to calculate the first ten principal components (PCs), summarising major axes of genetic variation across all samples. To investigate clustering within the native Australian dataset while minimising potential bias from drift in invasive populations, we performed discriminant analysis of principal components (DAPC) using the R package adegenet (Jombart, 2008). The optimal number of genetic clusters (K) was determined using the find.clusters function, exploring 1 ≤ K ≤ 15 and selecting the value of K associated with the lowest Bayesian Information Criterion (BIC). For the final DAPC, we retained three principal components and fitted K − 1 discriminant functions, visualising individuals along the first two discriminant axes.

#### 4.5.2 ADMIXTURE analysis

*ADMIXTURE* v. 1.3.0 was used to infer individual ancestry and population structure on the global dataset. Cross-validation was run for values of K = 1 to 10 and the optimal number of ancestral populations was determined by selecting the K with the lowest cross-validation error (i.e., K = 4; Figure S6).

#### 4.5.3 fineRADstructure analysis

To further investigate the population structure of *Ae. notoscriptus* and coancestry between individuals of the global dataset, we used the program *fineRADstructure*. The co-ancestry matrix of 191 individuals was based on a summary of nearest neighbour haplotype relationships, which were calculated using all the SNPs at each RAD locus. The individuals were grouped by similarity, and a phylogenetic tree was constructed using the *fineRADstructure* MCMC clustering algorithm with default settings. R scripts fineradstructureplot.r and finestructurelibrary.r (available at http://cichlid.gurdon.cam.ac.uk/fineRADstructure.html) were used for visualization.

#### 4.5.4 Treemix analysis

Using the seven population VCF genotype file, the occurrence of genetic drift was further investigated using *Treemix* v1.12. This software models the relationship among the sample groups and their ancestral groups using genome-wide allele frequency data and a Gaussian approximation of genetic drift (Pickrell and Pritchard 2012). We generated maximum-likelihood-based phylogenetic trees among all groups and iteratively, and we added two migration edges to the previously generated graph (*m*) (Fitak 2021). We rooted the graphs using the Noto3 samples as the outgroup based on analyses outlined in 3.5.1, 3.5.2 and 3.5.3. Additionally, a F3 test was run within *Treemix* (Reich et al. 2009) to test for admixture between Victorian Noto1, Noto2 and Noto3 populations based on signals observed in 3.5.3.

#### 4.5.5 Isolation by distance

We tested for isolation within the Victorian Noto1 only and Noto2 only datasets using the *mantel.randtest* function in the *R* (R Studio v1.4.1106 (R Core Team 2021)) package *ade4* (Dray and Dufour 2007). The simple Mantel tests analysed pairwise genetic distance matrices and the natural logarithm of Haversine pairwise geographic distance, employing 999 permutations and Bonferroni correction to assess statistical significance. Rousset’s *a* (Rousset 2000) provided genetic distances, calculated in *SPAGeDi* (Hardy and Vekemans 2002) for each group separately.

#### 4.5.6 Distance (d_XY_) tree inference

We used *pixy v2.0.0* (Korunes and Samuk 2021) to estimate pairwise genome-wide *d_XY_* among all samples. This used the VCF files processed in ***4.4.3.*** Genome-wide *d_XY_* estimates were bootstrapped 100 times by sampling 1 Mb windows with replacement and a minimum depth of 10. To visualize patterns of genetic differentiation, the resulting pairwise *d_XY_* matrix was used to construct an unrooted neighbor-joining tree in *R* (v4.3.2) using the package *ape* (Paradis et al. 2004). The *d_XY_* matrix was first converted into a distance object, and the neighbor-joining algorithm was applied to infer relationships among samples. The resulting tree was plotted and annotated in *R* to illustrate clustering of major genomic lineages.

#### 4.5.7 Autosomal heterozygosity and Tajima’s D

For individual autosomal heterozygosity, a VCF file for each individual was exported from the VCF files processed in ***4.4.3*** and filtered using bcftools v.1.16 to remove missing data sites, sites with rare alleles, sites with less than 15X coverage and sites with more than 100X coverage. Polyallelic sites were atomized into multiple SNPs using ‘bcftools norm -a --atom- overlaps’. For Tajima’s D, the same VCF files processed in ***4.4.3*** were used to calculate Tajima’s D in 100 kb windows using VCFtools, with each population filtered to retain only SNPs where every individual had a minimum depth of 10.

#### 4.5.8 Identification and characterisation of structural variants

We analysed patterns of population structure across the genome using the *R* package *lostruct* (Li and Ralph 2019) to detect regions of abnormal population structure that might be generated by structural variants using the Victorian Noto1 and Noto2 only datasets. The genome was divided into non-overlapping windows of 1000 SNPs and PCA was calculated for each window to reflect population structure. To measure the similarity of patterns of relatedness between windows, Euclidean distances were calculated using the first two PCs and these were then mapped in 40-dimensional space using multidimensional scaling (MDS). Different window sizes were tested to optimise the balance between signal and noise before a window size of 50 SNPs was chosen for all subsequent analyses.

Genomic regions with extreme MDS values were identified by defining outlier windows as those with absolute values greater than four standard deviations from the mean across all windows for each of the 40 MDS coordinates. The coordinates of outlier regions were then defined by the start position of the first outlier window and the end position of the last outlier window.

While inversions are a major driver of MDS outliers detected by *lostruct*, such outliers can also be generated by other processes, including linked selection. To distinguish between these possibilities, we performed additional analyses to test for population genomic signatures characteristic of inversions. Because recombination is suppressed within inversions, haplotype blocks with alternate orientations should evolve largely independently, leading to clear nucleotide differences between them. Consequently, for an inversion segregating within a population, a PCA of genetic variation is expected to separate individuals into three groups in the absence of lethal genes: the two alternative homokaryotypes and a heterokaryotype cluster.

To test for this pattern, we conducted PCAs using *SNPrelate* (Zheng et al. 2012), incorporating all SNPs from the putative inversion region. To identify the composition of groups of genotypes, we used the R function “kmeans” with the method developed by Hartigan and Wong 1979 to perform clustering on the first PC, using the maximum, minimum and middle of the range PC scores as the initial cluster centres. We subsequently calculated individual heterozygosity for each of the clusters using *PLINK*. Additionally, we calculated pairwise *F_ST_* between each of the clusters as well as for Victorian Noto1 and each of the two putative inversion karyotypes in Noto2 to identify which orientation may be shared with Victorian Noto1.

We identified candidate genes within the genomic outlier region on chromosome 2 as described in 3.5. Sequences corresponding to these outlier regions were then extracted from the *Ae. notoscriptus* reference assembly (as in 3.2.). To annotate potential functional elements, these sequences were queried against the *Ae. aegypti* reference genome (AaegL5) using *BLASTn* v. 2.14 (Altschul et al. 1990) with default parameters. For each query, the top hits were retained based on alignment length, percent identity and e-value.

From these BLAST results, we extracted the gene annotations associated with the highest-scoring hits, including gene identifiers, coordinates, and functional descriptions. This approach allowed us to generate a list of candidate genes located within the putative inversion region.

To investigate potential functional enrichment among these candidate genes, we performed a Gene Ontology (GO) analysis using *g:Profiler* (https://biit.cs.ut.ee/gprofiler/gost). Default settings were applied, including multiple testing correction using the Benjamini–Hochberg false discovery rate. Only protein-coding genes with valid annotations were included in the analysis. Enrichment was assessed across the three GO categories: molecular function, biological process, and cellular component. We then extracted significant GO terms (adjusted p-value < 0.05).

### 4.6 mtDNA preparation, sequencing, and phylogenetic analysis

We amplified the cytochrome oxidase subunit 1 (COI) gene (900 bp) of three to five samples per location (Table S6) using previously described primer combinations from Joy and Conn (2001) through PCR reactions. The PCR reactions were conducted in 50 µL volumes and followed an amplification profile consisting of an initial denaturation step at 94°C for 3 min, followed by 38 cycles of denaturation at 94°C for 1 min, annealing at 53°C for 1 min, and extension at 72°C for 1 min. A final extension step at 72°C for 5 min was also included. After amplification, we ran all samples on a 1% agarose gel to confirm successful amplification.

DNA sequences were generated using Applied Biosystem 3730 capillary analysers (Macrogen, Inc., Korea). We edited and aligned DNA sequences using *Geneious Pro* v5.5 (Drummond et al. 2011) and the *Clustal W* sequence alignment plug-in (Larkin et al. 2007). All available COI sequences of *Ae. notoscriptus* were downloaded from online databanks before we identified unique haplotypes (Table S6).

To generate the COI gene tree based on unique haplotypes only, we used Bayesian Inference (BI) methods from *MrBayes* 3.1.2 (Ronquist and Huelsenbeck 2003). Before BI analyses, *MrModeltest* (Nylander 2004) was used to identify the optimal nucleotide substitution model under the Akaike Information Criterion (AIC) (Akaike 1973). The BI analyses consisted of two parallel Metropolis-Coupled Markov Chain Monte Carlo (MCMCMC) runs with four chains each and a temperature setting of 0.02. A total of 10 million generations was run, with convergence of parallel runs indicated by an average standard deviation of split frequency values less than 0.01. Burnin and convergence for each run were determined by assessing the stabilization of the likelihood score using the software Tracer (Rambaut and Drummond 2007). Post-burnin trees were summarized as a 50% majority rule consensus tree with posterior probabilities as nodal support. All analyses started with a random starting tree and seed with no root specified. *Ae. aegypti* was used as an outgroup taxon to generate the tree.

To run species delineation analysis of the identified lineages, we built a Tamura-Nei Neighbour-Joining tree with no outgroup based on the COI sequence data. Samples in the tree were matched to the previous COI clade assigned and then the Geneious species delimitation plug-in (Masters et al. 2011) was used to generate statistics that estimate phylogenetic exclusivity of each clade, the probabilities of such exclusivity having arisen by chance in a random coalescent process and the degree to which the clades can be classified as species based on COI sequence data.

### 4.7 Wolbachia screening

Every individual used in this study was screened for the presence of the bacterium *Wolbachia*, using wsp81F and wsp691R primers (Braig et al. 1998) with the Roche LightCycler LC480 based on the protocol of (Lee et al. 2012).

## Declarations

### Ethics approval and consent to participate

Not applicable

### Consent for publication

Not applicable

### Availability of data and materials

All sequence data will be publicly accessible on the NCBI SRA through accession number XXX.

### Authors’ contributions

Conceptualization: VP, TLS, AAH; Funding acquisition: VP, AAH; Data curation, methodology, visualization, and formal analysis – population genomics: VP, TLS; Data curation, methodology, visualization, and formal analysis – reference genome: RR, GP, LNC; Investigation—field sample collection: VP, NMEH; Project administration and supervision: TLS, NMEH, AAH; Validation: TLS, AAH; Writing – original draft: VP; Writing – review and editing: VP, TLS, AAH, RR, GP, LNC, NMEH

## Acknowledgments

We thank Christopher Hardy for providing samples from the NT, QLD and NSW, Nathan McConnell, Emily Ferrill and Susanne Kluh for providing samples from the USA and Odwell Muzari for samples for Far North Queensland. We also thank Joshua Thia, Moshe Jasper, Kay Hodgins and Jonathan Wilson for helpful discussions and suggestions. For assistance with molecular work, we would like to thank Kelly Richardson and Katie Robinson.

