## Supplemental information for "Reproductive isolation and structural variation among sympatric lineages of an international mosquito pest"

### Supplementary material for: "Reproductive isolation and structural variation among sympatric lineages of an international mosquito pest"

Véronique Paris[1,2], Nancy M. Endersby-Harshman[1], Rahul Rane[3], Gunjan Pandey[4], Leon N. Court[4], Ary A. Hoffmann[1], Thomas L. Schmidt[5]

[1] Pest and Environmental Adaptation Research Group, School of BioSciences, Bio21 Institute, University of Melbourne, Parkville, Victoria, Australia

[2] Department of Microbiology and Immunology, Doherty Institute for Infection and Immunity, University of Melbourne, Melbourne, Victoria, Australia

[3] CSIRO Health and Biosecurity, Parkville, Victoria, Australia

[4] CSIRO Environment, Acton, Australian Capital Territory, Australia

[5] School of Life and Environmental Sciences, University of Sydney, Camperdown, New South Wales, Australia

**Table S1. Summary of genome sequencing quality metrics** and reference genome alignment statistics across *Aedes notoscriptus* lineages and sampling locations

| | | | **Proportion REF** | | **Missing data** | | **Depth of coverage** | |
|---|---|---|---|---|---|---|---|---|
| **Lineage** | **Location** | **n** | *Mean* | *SD* | *Mean* | *SD* | *Mean* | *SD* |
| | QLD | 3 | | | 0.049 | 0.015 | 14.30 | 1.92 |
| | NSW | 4 | | | 0.053 | 0.028 | 13.70 | 4.39 |
| Noto1 | NZ | 2 | 0.967 | 0.005 | 0.053 | 0.026 | 7.58 | 1.88 |
| | USA | 22 | | | 0.041 | 0.017 | 11.40 | 4.53 |
| | VIC | 122 | | | 0.038 | 0.042 | 14.90 | 6.84 |
| Noto2 | VIC | 33 | 0.849 | 0.025 | 0.059 | 0.032 | 12.20 | 6.02 |
| Noto3 | NT | 3 | 0.871 | 0.006 | 0.224 | 0.026 | 8.13 | 2.76 |

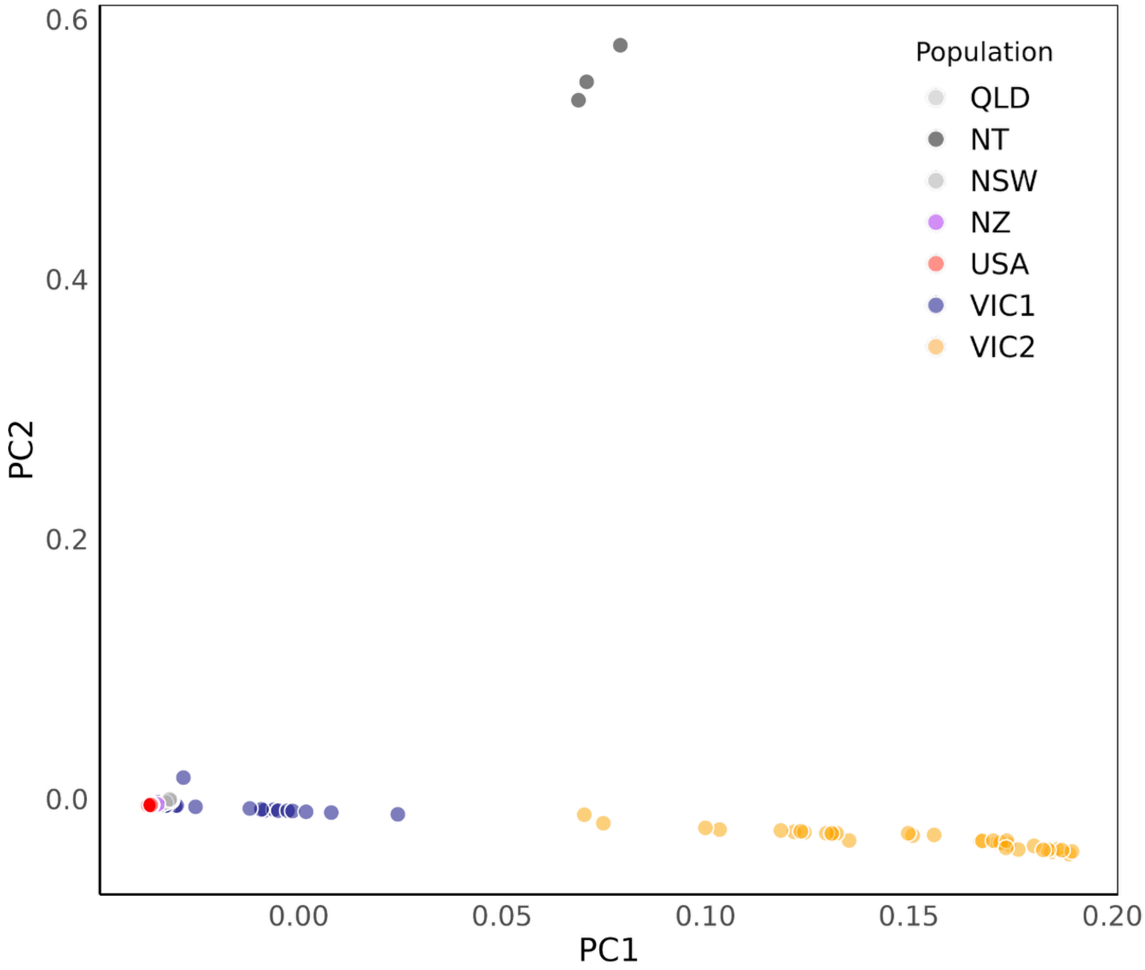


***Figure S1. Principal Component Analysis (PCA) of*** **Aedes notoscriptus** ***populations on the global SNP dataset***. *Each point represents an individual, coloured by population. PC1 and PC2 summarize the major axes of genetic variation. VIC1 individuals (blue) form a distinct cluster overlapping with other populations (NSW (green), QLD (light blue), NZ (grey), USA (red) representing Noto1, VIC2 (yellow) forms a separate cluster, representing Noto2, and NT (grey) is separated by PC2 from all other populations, representing Noto3.*

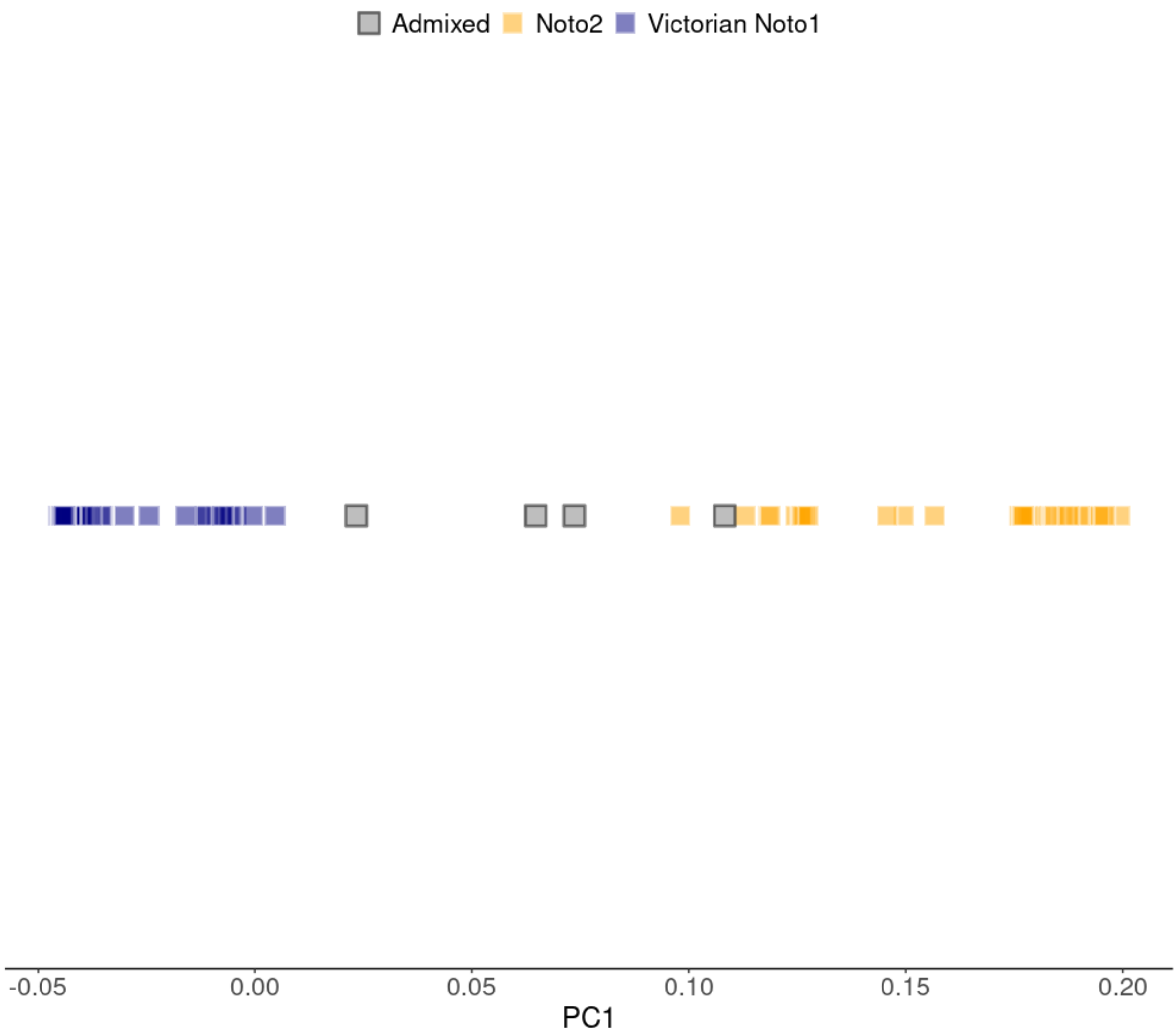


***Figure S2. Principal components analysis of the two Victorian populations and the putative admixed individuals identified by ADMIXTURE (Figure 3A).*** *Putative admixed individuals are projected onto a single principal component constructed from variation between non-admixed Victorian Noto1 and Noto2 individuals. The position of admixed individuals relative to the source populations reflects their admixture proportions.*

**COI sequencing results**

Partial sequences of the COI gene were obtained and genotyped for a total of 253 *Ae. notoscriptus* individuals. To enhance the analysis, all available COI sequences of *Ae. notoscriptus* from GenBank (Accession number: KF034536-KF034773) were downloaded, adding an additional 237 samples. This dataset yielded a total of 73 unique haplotypes, with the assignment of sample locations to haplotypes assigned to those of Endersby et al 2017 (Table S2).

Bayesian inference analyses and species delineation statistics conducted on the COI sequences revealed four highly supported monophyletic clades: red, blue, green, and brown (Figure S3). These colour codes follow Endersby et al. (2013), with the brown clade representing a newly identified lineage in this study. These clades exhibited strong posterior probabilities ranging from 0.95 to 1. Additionally, one clade showed moderate support (coded magenta, posterior probability 0.85), while another clade exhibited poor support (coded black, posterior probability 0.65) (Figure 3B inset and Figure S3). Detailed species delineation statistics are provided in Table S3.

The respective geographic distributions of the clades identified from the Bayesian inference phylogeny were largely consistent with those reported by Endersby et al. (2013). The blue clade appeared largely restricted to the subtropical regions of southern Queensland, with a few recordings in the tropical north of the state. The green clade was limited to tropical far north Queensland, while individuals belonging to the magenta and black clades were largely confined to temperate areas of southern Australia (VIC, southern NSW, and Western Australia), NZ, and the USA (black clade). The brown clade was restricted to the NT. In contrast, members of the red clade exhibited a broad distribution along the east coast of Australia, encompassing both tropical and temperate environments, and were also found in the USA. The phylogenetic network based on genome-wide $d_{xy}$ (Figure 2B) also highlighted mito-nuclear discordance, as the symbols and colours representing mitochondrial clades did not align perfectly with the nuclear genomic groupings (Figure 2B, insert).

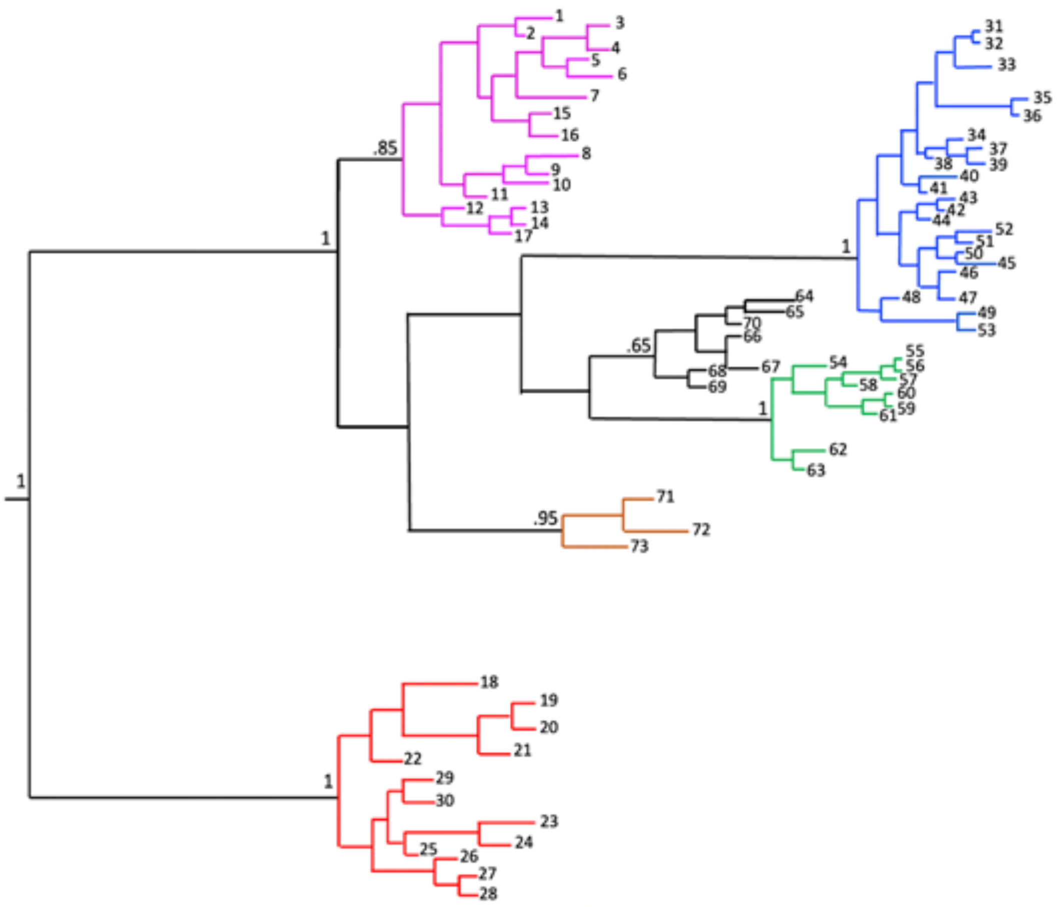


***Figure S3 Bayesian phylogeny of* Aedes notoscriptus.** *The analysis is based on 253 samples collected for this study as well as 237 sequences published by* Endersby et al. (2013)*. Clades are colour coded following* Endersby et al. (2013) *(red, magenta, blue, green and black), with the addition of the brown clade identified in this study. Membership of individuals to unique haplotypes can be found in Table S2.*

**Table S2 COI haplotypes of *Aedes notoscriptus* sorted by Bayesian phylogeny clade colour.**

| COI haplotype | Clade code | Sampling locations |
|---|---|---|
| 1 | Magenta | **VIC:** Tooradin (AJ008, AJ009) |
| 2 | Magenta | **VIC**: Port Melbourne (AN005, AN009, AN013), Bannockburn (V001), Inverleigh (U007), Dromana (P001), Mt Wallace (BD006), Craigieburn (AT004, AT005), Gembrook (AQ009, AQ010), Ferntree Gully (AR003), Cranbourne (AB004), Lonsdale (AD012), North Shore (AG009), Werribee (AH001), McKillops (K7, K5, K4, K1), Rye (L002), Romsey 1 6 7 11 17 (KF034654, KF034655, KF034656, KF034657, KF034658) |
| 3 | Magenta | **VIC:** Leopold (AF010), Bannockburn (V0007), Balliang (BD005), KF034633, KF034566 |
| 4 | Magenta | **VIC:** Balliang (BE003, BE012) |
| 5 | Magenta | **VIC:** Torquay (Q002), Bairnsdale (B4), Eltham (TT021) |
| 6 | Magenta | **VIC**: Mt Wallace (BD007), Healsville (BI012) |
| 7 | Magenta | **VIC:** Eynesbury (BG002, BG003, BG004, BG005, BG001), Baccus March (AX005) |
| 8 | Magenta | **VIC:** Wichelsea (T001, T006, T007, T008) |
| 9 | Magenta | **VIC**: St Andrews (BH001), Healesville (BI011) |
| 10 | Magenta | **VIC**: Mt Eliza (Z004), Rye (M012), Pearcedale (Y012) |
| 11 | Magenta | **VIC**: Flinders (R012), Leopold (AF013) |
| 12 | Magenta | **VIC**: Parkville (BA007), Albion (BB001, BB002) |
| 13 | Magenta | **VIC:** Bannockburn (V005) |
| 14 | Magenta | **WA:** WA 2 4 8 9 10 12 1Bb 3b 4b 6b 7b 8b 10b 2Bb (KF034687, KF034688, KF034689, KF034690, KF034691, KF034692, KF034695, KF034696, KF034697, KF034698, KF034699, KF034700, KF034701, KF034702) |
| 15 | Magenta | **VIC:** Warrandyte 6 9 (KF034680, KF034678) |
| 16 | Magenta | **VIC:** Broadford 12 17 (KF034564, KF034565), Lonsdale (AD014) |
| 17 | Magenta | **VIC**: Killops (K2, K3) |
| 18 | Red | **VIC:** Pakenham (AP006), Ferntree Gully (AR002, AR004, AR005, AR007), Winchelsea (T010), Bacchus March (AX011), Black Rock (AL006), Philip Island (AK001, AK003, AK010), Chelsea (AA006) |
| 19 | Red | **VIC:** Mt Eliza (Z013),<br>**QLD**: Mt Glorious 4 9 12 27 (KF034636, KF034637, KF034638, KF034639) |

| | | |
|---|---|---|
| 20 | Red | **VIC:** Tooradin (AJ013), Port Melbourne (AN010, AN011), Craigieburn (AT011), Balliang (BE011), Healesville (BI010), Broadford 9 (KF034569), Tecoma 10b 20b (KF034740, KF034741)<br>**NSW**: Georges River 27 (KF034588), c67, c66, c65, c73 |
| 21 | Red | **QLD:** James Cook Uni 7 8 9 (KF034605, KF034606, KF034607), Parramatta Park 18 19 (KF034645, KF034646) |
| 22 | Red | **VIC:** Montmorency 9 (KF034634), Tecoma 16b 22b (KF034746, KF034747)<br>**NSW:** Blue Mountain 7 (KF034563)<br>**QLD:** Kilcoy 5b 18b (KF034728, KF034729) |
| 23 | Red | **QLD**: Kilcoy 20b (KF034732) |
| 24 | Red | **QLD:** Mt Glorious 15 (KF034635) |
| 25 | Red | **VIC:** Warrandyte 7 (KF034679)<br>**NSW:** Black Town 3 (KF034553) Blue Mountain 11 (KF034558), Georges River 8 9 11 28 (KF034589, KF034590, KF034591, KF034592), Warriewood 15 21 (KF034681, KF034682), Hurstville (c56, c57, c60), Connells Point (c71).<br>**QLD:** Mt Isa 2 23 33 31 6 (KF034640, KF034641, KF034642, KF034643, KF034644) |
| 26 | Red | **VIC:** Bacchus March (AX004), Parkville (BA011) |
| 27 | Red | **VIC:** McKillops (K6)<br>**QLD:** Magnetic Island 33 (KF034612), Hervey Bay 5 (KF034596) |
| 28 | Red | **VIC**: Balliang (AE009), Leopold (AF004, AF008) |
| 29 | Red | **USA:** San Diego (D3) |
| 30 | Red | **USA:** San Diego (D4) |
| 31 | Blue | **QLD:** Tooheys Forest 7 (KF034665) |
| 32 | Blue | **QLD:** Tooheys Forest 14 (KF034660) |
| 33 | Blue | **QLD:** Cannon Hill 6 17 23 24 25 (KF034582, KF034581, KF034583, KF034584, KF034585) Hervey Bay 14 17 (KF034594, KF034595),<br>McDowall 23 24 (KF034625, KF034626), Mango Hill 21Bb (KF034627, KF034758) |
| 34 | Blue | QLD: Mareeba Nth 28 (KF034617), Seisia (S1, S5) |
| 35 | Blue | **QLD:** Buderim 21 (KF034570) |
| 36 | Blue | **QLD:** Mango Hill 11Bb (KF034760) |
| 37 | Blue | **QLD**: Mango Hill 14 Bb (KF034757) |
| 38 | Blue | **QLD:** Chinchilla 20b (KF034720) |

| | | |
|---|---|---|
| 39 | Blue | **QLD:** Jimboomba 48b (KF034770) |
| 40 | Blue | **QLD:** Charleville 11b (KF034726) |
| 41 | Blue | **QLD:** Kilcoy 15b (KF034727) |
| 42 | Blue | **QLD:** Mango Hill 16Bb (KF034762) |
| 43 | Blue | **QLD:** Chinchilla 20b (KF034719), Jimboomba 1Bb (KF034765) |
| 44 | Blue | **QLD:** Mango Hill 19Bb 20Bb (KF034763, KF034764) |
| 45 | Blue | **QLD:** Mango Hill 25Bb (KF034759) |
| 46 | Blue | **QLD:** Buderim 5 10 17 23 28 30 (KF034571, KF034572, KF034573, KF034574, KF034575, KF034576, KF034577) |
| 47 | Blue | **QLD:** Chinchilla 12b (KF034718), Jimboomba 3Bb 5b (KF034768, KF034769), Mango Hill 12Bb 15Bb (KF034755, KF034580), Cannon Hill 28 (KF034761), McDowall 21 (KF034622) |
| 48 | Blue | **QLD:** Chinchilla 34b (KF034716) |
| 49 | Blue | KF034661, KF034578, KF034599, KF034679, KF034662, KF034621, KF034600, KF034652, KF034647, KF034648, KF034711, KF034650, KF034756, KF034653, KF034713, KF03476, KF034651, KF034649, KF034663, KF034710, KF034712, KF034664, KF034730, KF034731 |
| 50 | Blue | **QLD:** Hervey Bay 5 (KF034596), McDowall 25 (KF034623), Charleville 6b 16b 19b 26b 34b (KF034721, KF034722, KF034723, KF034724, KF034725) |
| 51 | Blue | **QLD:** Jimboomba 9b (KF034767) |
| 52 | Blue | **QLD:** Tooheys Forest 9 16 (KF034666, KF034667) |
| 53 | Blue | **QLD**: Seisia (S3) |
| 54 | Green | **QLD:** Ayr population 1_1-6 1_1-12 (KF034539, KF034540) |
| 55 | Green | **QLD:** Ayr population 1_1-21 (KF034541) |
| 56 | Green | **NSW:** Black Town 11 (KF034549) |
| 57 | Green | **NSW:** Blue Mountain 28 (KF034562)<br>**QLD:** Chinchilla 1b (KF034717), Tooheys Forest 10 (KF034659), Wide Bay 6 (KF034714), Jimboomba 6Bb 7b 8b (KF034771, KF034772, KF034773) |
| 58 | Green | **QLD:** James Cook Uni 4 10 18 (KF034602, KF034603, KF034604) |
| 59 | Green | **QLD:** Mareeba Nth 6 18 29 (KF034618, KF034619, KF034620) |
| 60 | Green | **QLD:** Magnetic Island 11 16 20 23 (KF034608, KF034609, KF034610, KF034611) |

| | | |
|---|---|---|
| 61 | Green | **QLD:** Hervey Bay 12 (KF034593) Mareeba Nth 11 13 (KF034614, KF034615) |
| 62 | Green | **QLD:** Buderim 17 (KF034573), Ayr population 1_1-15 1_1-36, 2_2-3b 2_2-5b 2_2-8b 2_2_10b, 2_2_11b 2_2_13b 2_2_16b<br>(KF034536, KF034538, KF034542, KF034543, KF034544, KF034545, KF034546, KF034547, KF034548) |
| 63 | Green | **QLD**: Hervey Bay 6 8 (KF034597, KF034598), Magnetic Island 33 (KF034612), Mareeba Nth 14 (KF034616) |
| 64 | Black | **VIC:** Point Leo (W004, W007, W010, W014), Eltham (TT013, TT015, TT024), Torquay (Q005, Q007, Q010), Flinders (R001, R003, R009), Dromana (P005, P008), Ivanhoe (J002), Parkville (BA006), Tailors Lake (AV007), Philip Island (AK009), Black Rock (AL005), Mt Eliza (Z001), Crib Point (X008), North Shore (AG007, AG011), Little River (BF004), Pearcedale (Y003), Chelsea (AA009), Cranbourne (AB005), Narre warren (AC009), Healesville (BJ003), Montmorency 1 15 (KF034628, KF034629)<br>**NSW:** Black Town 27 (KF034552), Blue Mountain 17 22 (KF034559, KF03460), Hurstville (c63) |
| 65 | Black | **VIC:** Elwood (AM005), Sunbury (AW003, AW010), Tecoma 2b 4b 9b 11b 14b 15b (KF034749, KF034750, KF034751, KF034752, KF034753, KF034754), Bairnsdale (B1)<br>**NSW:** Black Town 1 (KF034551)<br>**WA:** WA 3 5 5b (KF034693, KF034694, KF034703) |
| 66 | **Black** | **VIC:** Inverleigh (U010), Mt Waverley (TT003, TT004), Eltham (TT018), Flinders (R010), Moriac (S004, S005), Dromana (P011), Ivanhoe (J007, J014), Albion (BB003), Sunbury (AW007, AW011, AW012), Bacchus March (AX007, AX012), Parkville (BA001), Tailors Lake (AV005, AV008, AV010), Mernda (AS009, AS010), Craigieburn (AT010), Black Rock (AL007), Gembrook (AQ001, AQ011), Tailors Lake (AV009), Philip Island (AK005), Crib point (X001), Pearcedale (Y009), Balliang (BE001), Mt. Eliza (Z011), Inverleigh (U003), Chelsea (AA001, AA005, AA010), Narre warren (AC002, AC005), Lonsdale (AD009, AD004), Rye (K001, M001, M004), Bairnsdale (B3, B5), Hoddles Creek (BJ002, BJ004), Montmorency 11 12 (KF034630, KF034631), Warrandyte 4 10 11 12 (KF034674, KF034675, KF034676, KF034677), Tecoma 1b 3b 5b 6b8b 17b19b (KF034733, KF034734, KF034735, KF034736, KF034737, KF034738, KF034739)<br>**US:** (G1, E1, D1, N5F, E2, J2, D2, N8F, J1, A2, H1, F1, A1)<br>**NSW:** Black Town 8 17 (KF034554, KF034555), Blue Mountain 10 30 (KF034556, KF034557), Georges River 12 26 (KF034586, KF034587), Upper Hutt 11 12 13 17 19 19 20 (KF034668, KF034669, KF034670, KF034671, KF034672, KF034673), Warriewood 3 8 24 29 (KF034683, KF034684, KF034685, KF034686), Hurstville (c58)<br>**ACT:** Palmerston (c54, c47, c52)<br>**WA:** WA 9b (KF034704),<br>**NZ:** Whangarei F1 F5 F7 O17 O21 (KF034705, KF034706, KF034707, KF034708, KF034709) |

| | | |
|---|---|---|
| 67 | **Black** | **VIC:** Pearcedale (Y002, Y010), Hoddles Creek (BJ005), Point Leo (W001), Crib Point (X010), Craigieburn (AT012), Cranbourne (AB003, AB010), Tecoma 7b 12b 13b 18b (KF034742, KF034743, KF034744, KF034745)<br>**NSW:** Georges River 26 (KF034587) |
| 68 | **Black** | **VIC:** Broadford 3 (KF034567), Bannockburn (V009), Moriac (S014), Mt Eliza (Z003), Narrewarren (AC011), Crib Point (X005) |
| 69 | **Black** | **USA:** San Diego (N6L1, D2, H1) |
| 70 | **Black** | **VIC:** Meredith (BC005, BC010, BC004, BC011, BC012), Black Rock (AL002), Dromana (P014), Tecoma 21b (KF034748), Cranbourne (AB006), North Shore (AG008), Montmorency 3 (KF034632)<br>**NSW:** Black Town 12 (KF034550), Blue Mountain 18 (KF034561)<br>**ACT:** Palmerston (c50) |
| 71 | **Brown** | **NT:** Darwin (c1) |
| 72 | **Brown** | **NT:** Darwin (c3, c4) |
| 73 | **Brown** | **NT:** Darwin (c2, c5) |

**Table S3 Species delineation statistics for COI sequence data from *Aedes notoscriptus*.**

| Clade code | Closest Species | Mono-phyletic | Intra dist[1] | Inter dist – closest[2] | Intra/ Inter | *P* ID (strict)[3] | *P* ID (liberal)[4] | *P* (randomly distinct)[5] | Rosenberg's *P* (AB)[6] |
|---|---|---|---|---|---|---|---|---|---|
| Blue | Black | Yes | 0.005 | 0.028 | 0.16 | 0.94 | 0.980 | <0.05 | 9.5E-11 |
| Red | Blue | Yes | 0.007 | 0.053 | 0.14 | 0.90 | 0.97 | <0.05 | 9.5E-11 |
| Green | Black | Yes | 0.008 | 0.012 | 0.67 | 0.74 | 0.92 | 1 | 7.6E-9 |
| Black | Green | No | 0.004 | 0.012 | 0.37 | 0.85 | 0.95 | 0.33 | NA |
| Magenta | Black | No | 0.004 | 0.015 | 0.280 | 0.90 | 0.97 | 0.17 | NA |
| Brown | Magenta | Yes | 0.001 | 0.020 | 0.07 | 0.82 | 0.97 | <0.5 | 4.4E-9 |

[1] Intra Dist, the average pairwise tree distance among members of a putative species. [2] Inter Dist, the average pairwise tree distance between the members of one putative species and the members of the closest second putative species. [3] *P* ID (strict), probability of correctly identifying an unknown member of the putative species using the criterion that must fall within, but not sister to, species clade in a tree. [4] *P* ID (liberal), the mean probability of correctly identifying an unknown member of the putative species using the criterion that it falls within, or sister to, the species clade in a tree. [5] *P* (randomly distinct), probability that a clade had the observed degree of distinctiveness due to random coalescent processes. [6] Rosenberg's *P* (AB), probability of reciprocal monophyly under a random coalescent model.

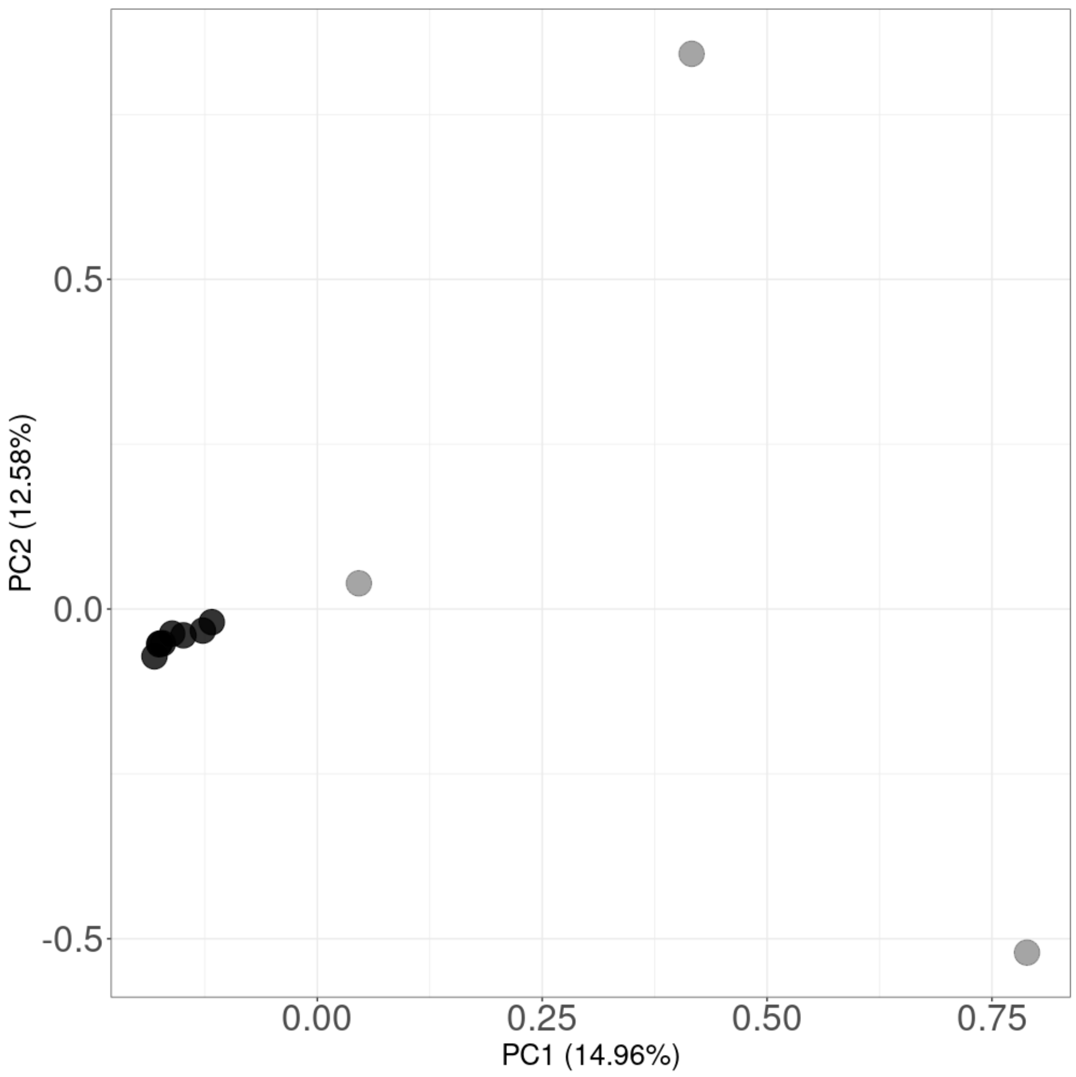

***Figure S4 Principal component analysis of putative admixed individuals between Victorian Noto1 and Noto2.*** *Black circles show individuals identified through fineRADstructure while grey circles show individuals identified through ADMIXTURE.*

**Table S4 Observed counts of Victorian Noto1 and Noto2 individuals at sites where both lineages co-occur across Greater Melbourne**, together with lineage frequencies (p and q) and the expected number of hybrids under random mating (2pq × Total) at each location.

| Location | Victorian Noto1 | Noto2 | Total | p = VIC1/Total | q = VIC2/Total | Expected hybrids 2pq*Total |
|---|---|---|---|---|---|---|
| Tooradin | 2 | 2 | 4 | 0.5 | 0.5 | 2 |
| Port Melbourne | 4 | 1 | 5 | 0.8 | 0.2 | 1.6 ≈ 2 |
| Gembrook | 3 | 2 | 5 | 0.6 | 0.4 | 2.4 ≈ 2 |
| Meredith | 1 | 2 | 3 | 0.333 | 0.667 | 1.33 ≈ 1 |
| Balliang | 1 | 3 | 4 | 0.25 | 0.75 | 1.5 ≈ 2 |
| St. Andrews | 1 | 4 | 5 | 0.2 | 0.8 | 1.6 ≈ 2 |
| Healesville | 1 | 3 | 4 | 0.25 | 0.75 | 1.5 ≈ 2 |
| Winchelsea | 1 | 4 | 5 | 0.2 | 0.8 | 1.6 ≈ 2 |
| Inverleigh | 3 | 2 | 5 | 0.6 | 0.4 | 2.4 ≈ 2 |
| Bannockburn | 2 | 3 | 5 | 0.4 | 0.6 | 2.4 ≈ 2 |
| Point Leo | 4 | 1 | 5 | 0.8 | 0.2 | 1.6 ≈ 2 |

**Table S5 Top ten genes identified within the chromosome 2 outlier region based on cumulative BLAST hit counts against the *Aedes aegypti* reference genome.** Total BLAST hits represent the number of sequences matches per gene across all detected transcript variants; the number of transcript variants indicates the count of distinct annotated isoforms contributing to these hits.

| Gene ID | Product description | Total BLAST hits | No. transcript variants |
|---|---|---|---|
| LOC5569057 | Attractin | 4,655 | 2 |
| LOC5564358 | Serine/threonine-protein kinase 32A | 3,610 | 9 |
| LOC110676084 | Uncharacterized protein | 3,163 | 1 |
| LOC5568545 | Calpain-D | 2,038 | 4 |
| LOC5564718 | Zwei Ig domain protein zig-8 | 1,867 | 2 |
| LOC5574477 | Tyrosine-protein kinase receptor | 1,508 | 7 |
| LOC5573523 | Sterol regulatory element-binding protein | 1,410 | 1 |
| LOC5577895 | Down syndrome cell adhesion molecule-like protein | 1,393 | 8 |
| LOC5577573 | Uncharacterized protein | 1,363 | 2 |

| | | | |
|---|---|---|---|
| LOC5569913 | Serine-rich adhesin for platelets | 1,269 | 2 |

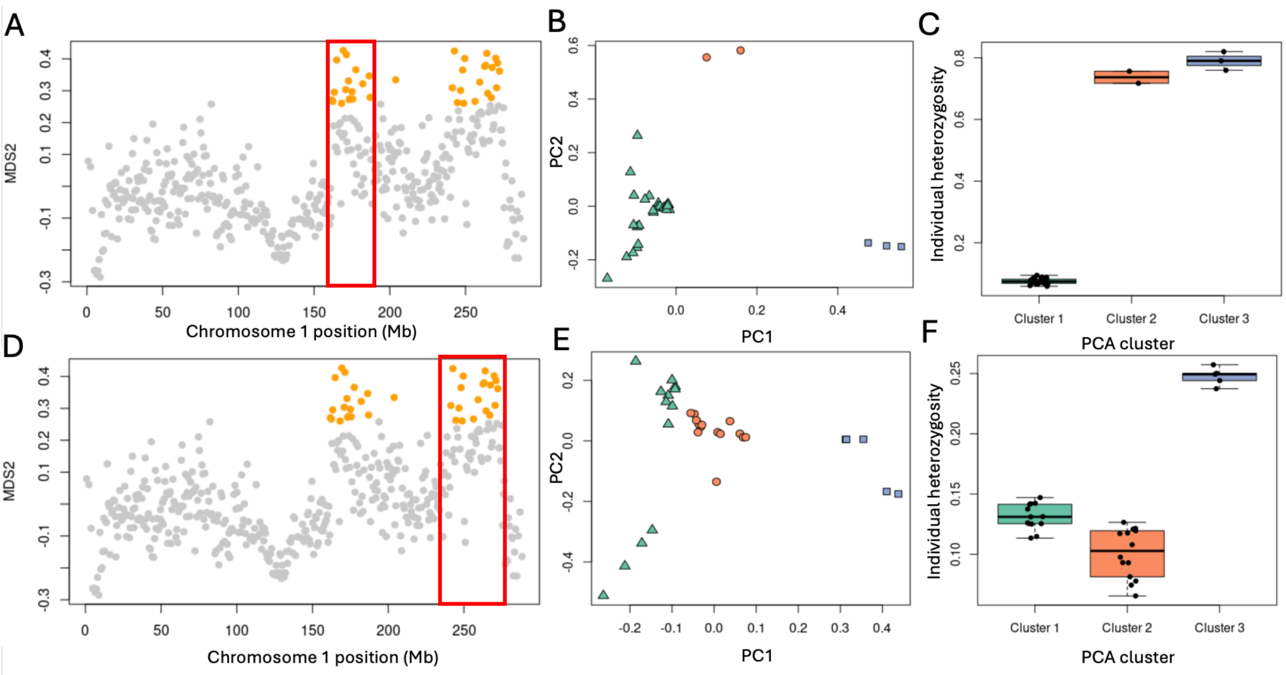


***Figure S5 Characterisations of the MDS outlier region on chromosome 1 of Noto2 lineage.*** *(A) + (D) Plot of MDS values across chromosome 1. Each grey dot represents a window of 50 SNPs and outlier windows are highlighted in red. (B) + (E) PCA based on SNPs from outlier region. Three clusters identified and illustrated in green triangles, orange circles and blue squares. (C) + (F) Heterozygosity for each of the groups identified in PCA*

**Table S6 Sampling locations and number individuals included in the genomic, mtDNA and *Wolbachia* analysis.**

| Location | Latitude | Longitude | Provided by | # ddRADseq | # COI | # Wolbachia screened |
|---|---|---|---|---|---|---|
| **Victoria** | | | | | | |
| Mt Eliza | -38.1914 | 145.0858 | VP | 4 | 5 | 5 |
| Chelsea | -38.0479 | 145.1153 | VP | 5 | 5 | 5 |
| Black Rock | -37.9843 | 145.0248 | VP | 3 | 4 | 4 |
| Elwood | -37.8843 | 144.9803 | VP | 1 | 1 | 1 |
| Cranbourne | -38.0995 | 145.2687 | VP | 5 | 5 | 5 |
| Narre Warren | -37.9840 | 145.3160 | VP | 2 | 5 | 5 |
| Pakenham | -38.0857 | 145.4776 | VP | 1 | 1 | 1 |
| Torquay | -38.3278 | 144.3201 | VP | 5 | 5 | 5 |
| Moriac | -38.2489 | 144.1704 | VP | 2 | 5 | 5 |
| Portarlington | -38.1215 | 144.6582 | VP | 3 | 5 | 5 |

| Location | Latitude | Longitude | Provided by | # ddRADseq | # COI | # Wolbachia screened |
|---|---|---|---|---|---|---|
| Werribee | -37.9205 | 144.7016 | VP | 2 | 3 | 3 |
| Altona | -37.8721 | 144.8168 | VP | 5 | 5 | 5 |
| Albion | -37.7783 | 144.8206 | VP | 4 | 4 | 4 |
| Little River | -37.9626 | 144.5067 | VP | 1 | 1 | 1 |
| Phillip Island | -38.4524 | 145.2477 | VP | 4 | 5 | 5 |
| Inverleigh | -38.1041 | 144.0588 | VP | 5 | 5 | 5 |
| Gembrook | -37.9543 | 145.5508 | VP | 5 | 5 | 5 |
| Ferntree Gully | -37.8801 | 145.2669 | VP | 4 | 5 | 5 |
| Mt Waverley | -37.8828 | 145.1250 | VP | 2 | 2 | 2 |
| Lilydale | -37.7551 | 145.3528 | VP | 4 | 4 | 4 |
| Taylors Lakes | -37.7071 | 144.8005 | VP | 4 | 5 | 5 |
| Bacchus Marsh | -37.6723 | 144.4417 | VP | 4 | 5 | 5 |
| Mt Wallace | -37.7476 | 144.2340 | VP | 1 | 2 | 2 |
| Ivanhoe | -37.7731 | 145.0463 | VP | 5 | 5 | 5 |
| Parkville | -37.7819 | 144.9563 | VP | 5 | 5 | 5 |
| Dromana | -38.3373 | 144.9621 | VP | 5 | 5 | 5 |
| Rye | -38.3732 | 144.7912 | VP | 0 | 5 | 5 |
| Hoddles Creek | -37.8173 | 145.5756 | VP | 5 | 5 | 5 |
| Flinders | -38.4686 | 145.0225 | VP | 5 | 5 | 5 |
| Point Leo | -38.4191 | 145.0758 | VP | 5 | 5 | 5 |
| Winchelsea | -38.2449 | 143.9950 | VP | 5 | 5 | 5 |
| Bannockburn | -38.0490 | 144.1680 | VP | 5 | 5 | 5 |
| Crib Point | -38.3612 | 145.2023 | VP | 5 | 3 | 5 |
| Pearcedale | -38.2070 | 145.2347 | VP | 5 | 5 | 5 |
| Tooradin | -38.2140 | 145.3792 | VP | 4 | 5 | 5 |
| Port Melbourne | -37.8414 | 144.9361 | VP | 5 | 5 | 5 |
| Eynesbury | -37.7930 | 144.5604 | VP | 5 | 5 | 5 |
| Meredith | -37.8422 | 144.0751 | VP | 3 | 5 | 5 |
| St Andrews | -37.5989 | 145.2774 | VP | 5 | 5 | 5 |
| Balliang | -37.8735 | 144.3810 | VP | 4 | 5 | 5 |
| Healesville | -37.6587 | 145.5114 | VP | 4 | 4 | 4 |
| Longsdale | -38.2691 | 144.6140 | VP | 0 | 5 | 5 |
| Craigieburn | -37.5900 | 144.9300 | VP | 0 | 4 | 4 |
| Mernda | -37.5900 | 145.1000 | VP | 0 | 2 | 2 |
| Eltham | -37.7146 | 145.1568 | VP | 3 | 5 | 5 |
| Sunbury | -37.5861 | 144.7268 | VP | 0 | 5 | 5 |

| Location | Latitude | Longitude | Provided by | # ddRADseq | # COI | # Wolbachia screened |
|---|---|---|---|---|---|---|
| Leopold | -38.1832 | 144.4786 | VP | 0 | 5 | 5 |
| North Shore | -38.1005 | 144.3733 | VP | 0 | 5 | 5 |
| **East Gippsland** | | | | | | |
| Bairnsdale | -37.8404 | 147.5893 | VP | 0 | 5 | 5 |
| McKillops Bridge | -37.0915 | 148.4135 | VP | 0 | 5 | 5 |
| **New South Wales** | | | | | | |
| Connells Point | -33.9839 | 151.0954 | Christopher Hardy | 1 | 5 | 5 |
| Hurstville | -33.9675 | 151.0972 | Christopher Hardy | 1 | 5 | 5 |
| **Australian Capital Territory** | | | | | | |
| Palmerston | -35.1938 | 149.1190 | Christopher Hardy | 2 | 5 | 5 |
| **Queensland** | | | | | | |
| Seisia | -10.8519 | 142.3719 | Odwell Muzari | 0 | 4 | 4 |
| Brisbane | -27.4776 | 153.0313 | Christopher Hardy | 3 | 5 | 5 |
| **Northern Territory** | | | | | | |
| Darwin | -12.4531 | 130.8440 | Tomoko Okazaki | 3 | 4 | 4 |
| **New Zealand** | | | | | | |
| Auckland | -36.9080 | 174.6307 | Rachel Cane | 2 | 0 | 3 |
| **USA** | | | | | | |
| San Diego | 32.7524 | -116.9744 | N. McConnell, E. Ferrill, S. Kluh | 22 | 5 | 20 |

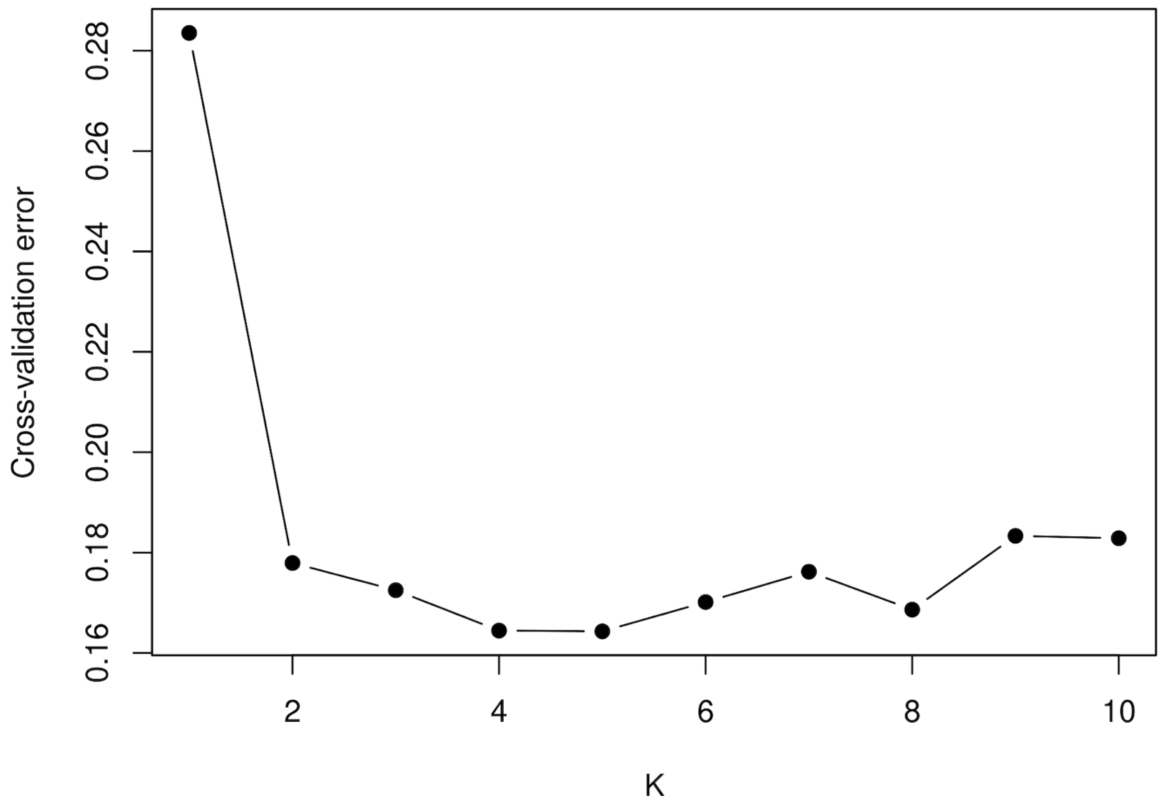


***Figure S6 ADMIXTURE cross-validation error for values of K = 1–10***. *Cross-validation error decreased sharply at low K and reached its minimum at K = 4, indicating that a four-cluster model best captures the underlying population structure in the global dataset. This value was therefore used for the main ADMIXTURE results presented in Figure 2*
